# A systematic quantitative approach comprehensively defines domain-specific functional pathways linked to *Schizosaccharomyces pombe* heterochromatin regulation

**DOI:** 10.1101/2024.02.13.579970

**Authors:** Abubakar Muhammad, Zsuzsa Sarkadi, Thomas van Emden, Agnisrota Mazumder, Matias Capella, Gergely Fekete, Vishnu N. Suma Sreechakram, Bassem Al-Sady, Balázs Papp, Ramón Ramos Barrales, Sigurd Braun

## Abstract

Heterochromatin plays a critical role in regulating gene expression and maintaining genome integrity. While structural and enzymatic components have been linked to heterochromatin establishment, a comprehensive view of the underlying pathways at diverse heterochromatin domains remains elusive. Here, we developed a systematic approach to identify factors involved in heterochromatin silencing at pericentromeres, subtelomeres, and the silent mating type locus in *Schizosaccharomyces pombe*. Using quantitative measures, iterative genetic screening, and domain-specific heterochromatin reporters, we identified 369 mutants with different degrees of reduced or enhanced silencing. As expected, mutations in the core heterochromatin machinery globally decreased silencing. However, most other mutants exhibited distinct qualitative and quantitative profiles that indicate domain-specific functions. For example, decreased mating type silencing was linked to mutations in heterochromatin maintenance genes, while compromised subtelomere silencing was associated with metabolic pathways. Furthermore, similar phenotypic profiles revealed shared functions for subunits within complexes. We also discovered that the uncharacterized protein Dhm2 plays a crucial role in maintaining constitutive and facultative heterochromatin, while its absence caused phenotypes akin to DNA replication-deficient mutants. Collectively, our systematic approach unveiled a landscape of domain-specific heterochromatin regulators controlling distinct states and identified Dhm2 as a previously unknown factor linked to heterochromatin inheritance and replication fidelity.

---

Heterochromatin, a fundamental form of DNA packaging found across eukaryotic genomes, plays pivotal roles in regulating gene expression, maintaining genome stability, shaping chromosomal architecture, and determining cell identity. Heterochromatin regions are associated with transcriptionally repressed chromatin and are characterized by a condensed structure, low histone acetylation, and the accumulation of specific histone modifications, notably methylation of lysine 9 of histone H3 (H3K9me). This histone mark is recognized by chromodomain proteins and self-propagated through a ‘‘read-write’’ mechanism (Allshire and Madhani, 2018; Grewal, 2023). Heterochromatin assembly comprises distinct steps: nucleation, spreading, and maintenance. Nucleation involves the DNA- or RNA-guided recruitment of a histone methyltransferase often involving multiple cycles that amplify the initial signal (Holoch and Moazed, 2015). Spreading describes the sequence-independent expansion of H3K9me-marked heterochromatin along the chromosome. Maintenance involves the stable formation of heterochromatin domains and their inheritance during DNA replication. Such inheritance depends on the self-templated propagation of repressive histone modifications as new nucleosomes assemble (Allshire and Madhani, 2018; Grewal, 2023). While constitutive heterochromatin persists throughout the cell cycle, often at gene-poor repetitive sequences, facultative heterochromatin forms on specific developmental or lineage-specific genes to stabilize distinct cell states. The spatial positioning of heterochromatin at the nuclear periphery further facilitates its assembly and maintenance (Grewal, 2023; Harr et al., 2016).

The fission yeast, *S. pombe*, has distinct constitutive heterochromatin domains present at the pericentromeric repeats, subtelomeres, and the silent mating-type locus (Allshire and Ekwall, 2015). Many conserved factors in metazoan heterochromatin assembly have orthologs in *S. pombe*, which are encoded by single-copy genes, offering a model system with reduced redundancy and complexity. For example, the homolog of Su(var)3-9, Clr4, is the sole H3K9 methyltransferase in *S. pombe* and catalyzes mono-, di-, and trimethylation (Bannister et al., 2001; Nakayama et al., 2001). Clr4 associates with a multimeric ubiquitin ligase to form CLRC (Hong et al., 2005; Horn et al., 2005; Jia et al., 2005; Li et al., 2005; Thon et al., 2005), which mediates H3K14 ubiquitylation, a prerequisite for H3K9 methylation (Oya et al., 2019; Stirpe et al., 2021). Heterochromatin assembly is initiated by CLRC recruitment to nucleation sites through DNA- and RNA-guided processes. This step involves DNA-binding factors (Cooper et al., 1997; Jia et al., 2004; Kanoh et al., 2005) and the RNA interference (RNAi) machinery including the argonaute-containing RNA-induced transcriptional silencing complex (RITS) and additional components (Bayne et al., 2010; Hayashi et al., 2012; Motamedi et al., 2004; Noma et al., 2004; Rougemaille et al., 2012; Sugiyama et al., 2005; Verdel et al., 2004). While RNAi is indispensable for heterochromatin establishment and maintenance at pericentromeres, it acts redundantly with DNA-binding factors at subtelomeres and the mating-type locus (Hansen et al., 2006; Jia et al., 2004). CLRC can also be recruited independently of RNAi to facultative heterochromatin via RNA-elimination factors or components of the telomere-protecting shelterin complex (Egan et al., 2014; Lee et al., 2013; Tashiro et al., 2013; Zofall et al., 2016, 2012).

Upon deposition, H3K9me recruits HP1 homologs (Swi6 and Chp2) and Clr4 itself, establishing a heterochromatic platform that governs the recruitment of additional heterochromatin factors. Among those factors is the Snf2-like nucleosome remodeler and histone-deacetylase repressor complex SHREC, an ortholog of mammalian NuRD (Job et al., 2016; Motamedi et al., 2008; Sugiyama et al., 2007). HP1 also acts as a docking site for Epe1, a putative H3K9me demethylase that counteracts heterochromatin spreading (Ayoub et al., 2003; Zofall and Grewal, 2006). Epe1 distribution on chromatin is confined to the heterochromatin boundaries through selective degradation by the ubiquitin ligase Cul4-Ddb1^Cdt2^, adding another layer of regulation (Braun et al., 2011). The association of HP1-bound factors is further controlled by phosphorylation and interaction with inner nuclear membrane proteins (Barrales et al., 2016; Holla et al., 2020; Shimada et al., 2009; Shipkovenska et al., 2020). In several systems, heterochromatin assembly is also subject to metabolic regulation and dependent on nutrient availability, such as methionine that serves as a donor precursor for histone methylation (Fan et al., 2015; Mentch et al., 2015; Serefidou et al., 2019). However, the broader spatio-temporal regulation of heterochromatin and the distinct requirements across different heterochromatin domains remain largely unexplored, underscoring a significant gap in our understanding of heterochromatin biology.

Genetic screens combined with reporter genes that monitor the transcriptional activity in individual heterochromatin domains *in vivo* have emerged as powerful tools for identifying heterochromatin regulators (Allshire and Ekwall, 2015). However, previous studies focused on single heterochromatin loci relied on qualitative color-based readouts or semi-quantitative methods with limited resolution. To overcome these limitations, here we adopt a quantitative and systematic approach targeting all major constitutive heterochromatin domains. By quantifying the degree of de-repression in these domains, we systematically determined the requirements for specific heterochromatin regulators. Moreover, by correlating phenotypic profiles across distinct heterochromatin domains, we uncovered striking phenotypic similarities among chromatin regulators belonging to the same complex, suggesting the potential to predict novel functional relationships. Additionally, we identified various metabolic pathway genes specifically required for subtelomeric silencing. Further, we identified and characterized a novel heterochromatin regulator, Dhm2, required for constitutive and facultative heterochromatin maintenance and connected to DNA replication. Our findings yield a substantial body of knowledge that will pave the way for future investigations into heterochromatin regulation.

## Results

### A quantitative and systematic screening approach to identify regulatory factors for all major constitutive heterochromatin domains

To systematically identify factors that regulate constitutive heterochromatin, we used a quantitative reporter assay to screen the *S. pombe* haploid deletion collection for altered heterochromatic silencing. Reporter strains carried the *ura4*^*+*^ gene at the pericentromeres (left innermost repeats of centromere 1, *imr1L; CEN*), the silent mating type locus (downstream of *mat3M; MAT*), the subtelomeres (7 kb downstream of the telomeric repeats of the right arm of chromosome 2; *SUBTEL*), or next to the telomeric repeats (left arm of chromosome 2; *TEL*) (Suppl. Figure S1a) (Allshire et al., 1995; Ekwall et al., 1999; Kanoh et al., 2005; Nimmo et al., 1998). A hygromycin resistance marker (*hphMX6*) inserted in euchromatin next to the heterochromatin locus allowed selection of the reporter (Suppl. Figure S1b). These reporter strains were crossed with a *kanMX6*-marked collection of 2,988 deletion mutants derived from a commercial mutant library of non-essential genes (Bioneer v3) that omits several mitochondrial protein-encoding genes and severely sick mutants (Suppl. Table S1). Large-scale genetic crosses were performed following high-throughput SGA (synthetic gene array) approaches (Baryshnikova et al., 2010; Verrier et al., 2015). To monitor the level of *ura4*^*+*^ silencing after crosses, we quantitatively measured gain or loss of growth on solid media lacking uracil (-URA) and containing 5-fluoroorotic acid (+FOA), which is converted into a toxic compound by the *ura4*^*+*^ gene product, respectively. We normalized these values to growth on non-selective media (relative growth) to account for pleiotropic effects affecting the overall fitness of the mutants and position effects from neighboring mutants (Barrales et al., 2016). Thus, changes in relative growth are proportional to the degree by which heterochromatin is compromised but independent of any other parameters, allowing the quantitative assessment of changes in heterochromatin states in the individual mutants in a highly reproducible manner (Suppl. Figure S1c).

For every reporter, we conducted multiple independent screens using the mutant collection (n = 3-8; Suppl. Tables S2, S3) and applied a multistep data processing pipeline to identify mutants significantly affecting heterochromatin silencing (Figure 1a; for details, see Methods). In brief, we applied z-normalization and median-centering to scale the relative growth data, allowing comparison across the four different reporter screens. For each biological replicate, we then combined the normalized values from the two readouts (-URA, +FOA), which we refer to as the *combined FOA/URA score*. This step enhanced the sensitivity of readouts and helped overcome limitations where certain mutants exhibited mild effects in one readout (e.g., +FOA) but substantial effects in the other (e.g., -URA). For high-confidence identification of candidates with altered silencing, we applied a cut-off using a P-value < 0.05 and an effect size threshold for the combined +FOA/-URA scores > 2.5 (for *CEN* we applied a threshold > 3 because of the leaky repression of the *imr1L* locus). To assess the validity of these parameters, we evaluated their ability to retrieve known heterochromatin factors employing common GO (Gene Ontology) terms (recall; Suppl. Table S4; Suppl. Figure S2). In addition, we analyzed the precision of our selection criteria by measuring the transcript levels of the *ura4*^*+*^ reporter gene for a representative subset of mutants (positive predicted value; Suppl. Table S5; Suppl. Figure S3). Overall, we find a good agreement between reporter growth data and transcript levels (Suppl. Table S6), validating the ability of our growth-based reporter approach to quantitate heterochromatin silencing.

**Figure 1:**
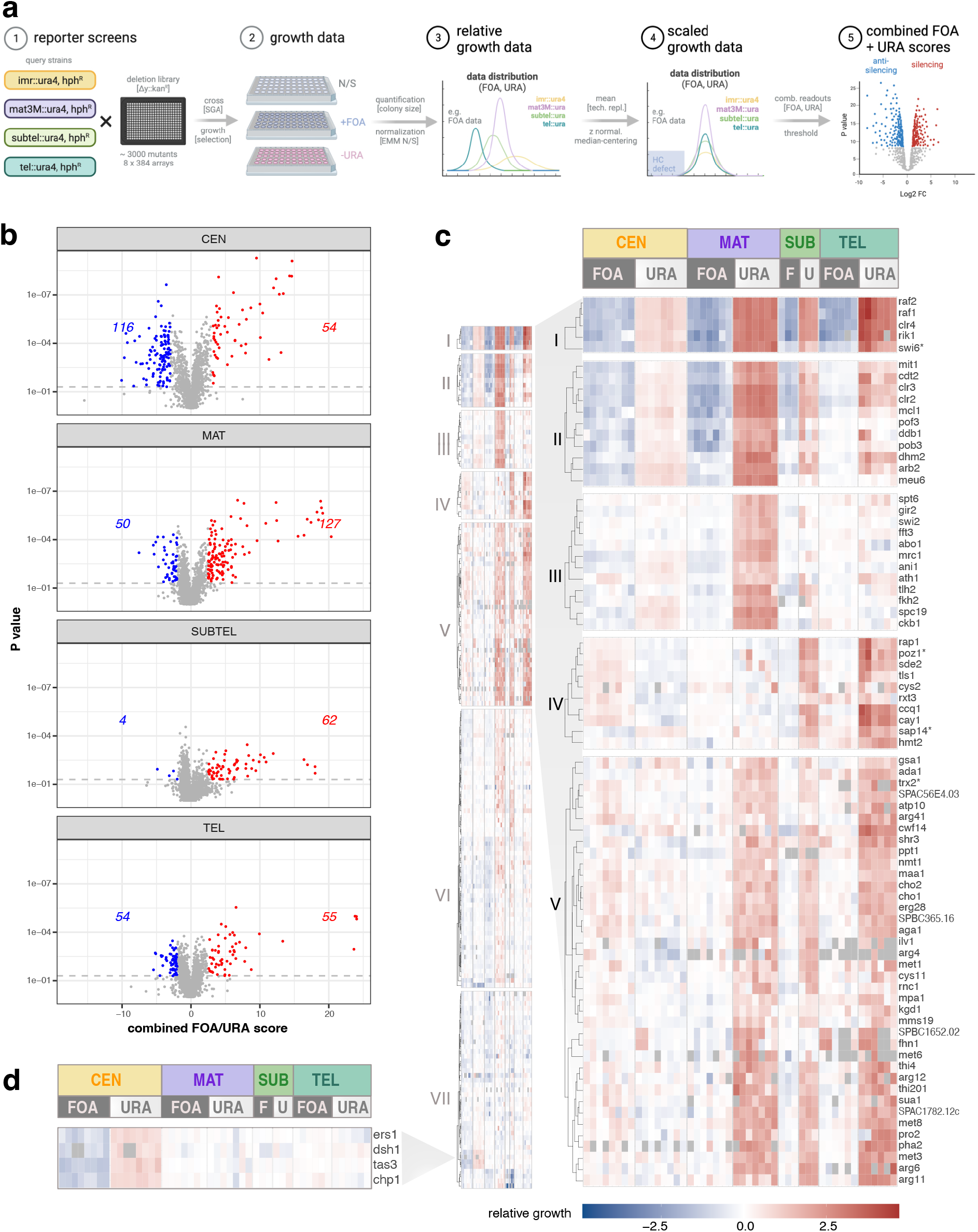
Systematic identification of heterochromatin regulators. **a)** Schematic overview of the screening strategy employing high-throughput silencing assays. The *ura4*^*+*^ gene in the reporter strains was positioned at the pericentromeric region (*imr1L; CEN*), the silent mating type locus (*mat3M; MAT*), the subtelomeric region (*subtel2R; SUBTEL*), or adjacent to telomeric repeats (*tel2L; TEL*). An additional hygromycin selection (*hphR)* marker was introduced adjacent to the heterochromatic domains. **b**) Volcano plots illustrating the combined FOA/URA scores (x-axis) and p-values (y-axis) derived from one-sample Student’s t-test for each mutant (details in Methods). The data include 3-8 independent biological replicates (*CEN*: 8; *MAT*: 7; *SUBTEL*: 3; *TEL*: 6), each comprising 2-4 technical replicates. Mutants with significantly altered silencing (p<0.05) are color-coded: red for silencing factors and blue for anti-silencing factors, with the respective number indicated. **c)** Heatmaps displaying k-means cluster analysis of log2-transformed relative growth values (+FOA, -URA) from 176 silencing mutants exhibiting a significant decrease in heterochromatin silencing in at least one domain (details in text). The left panel displays an overview of all clusters, while the right panel shows a subset of clusters (I-V) with gene names. The gene order within each cluster was determined through subsequent hierarchical clustering. **d)** Heatmap showing relative growth values of a subset of genes involved in RNAi from Cluster VII. In cases where mutants were misannotated in the gene deletion collection, the correct gene name is indicated by an asterisk (see Suppl. Table S8 and Methods).

### Identification of factors promoting and antagonizing heterochromatin silencing

By applying these selection criteria, we identified 180 genes that significantly reduced heterochromatic silencing of the *ura4*^*+*^ reporter to various extents. While several mutants affected silencing at multiple heterochromatin domains, the number of hits varied for each reporter locus. The screens performed at *MAT* retrieved the largest number of candidates (127 genes); fewer mutants affected silencing at *CEN* (54), *SUBTEL* (62), and *TEL* (55) (Figure 1b; Suppl. Table S3). We validated the screen by assessing endogenous heterochromatic transcripts by reverse transcriptase combined with quantitative PCR (RT-qPCR) for a representative selection of mutants. For 42 out of 53 candidates examined (Suppl. Table S7), we detected elevated levels of these transcripts (> 1.5-fold relative to WT), largely confirming the growth-based silencing defects in these mutants. The top hits included known members of the core heterochromatin machinery, namely CLRC (Clr4, Rik1, Raf1, and Raf2), SHREC (Clr3, Clr2, and Mit1), and the HP1 homolog Swi6 (Suppl. Figure S4a). These mutants showed highly reproducible values, underscoring the robustness and reproducibility of the quantitative readout and data normalization. In addition, they often displayed reciprocal readouts for the two reporter growth conditions (+FOA, -URA; Suppl. Figure S4a, b). Factors involved in RNAi (Chp1, Tas3, Dsh1, Ers1) were exclusively detected by the *imr1L::ura4* reporter (*CEN*), consistent with their essential function at pericentromeres but redundant role at the other heterochromatin regions. Of note, our screen did not identify certain RNAi components (Ago1, Dcr1, Rdp1, Hrr1) and known heterochromatin factors (Sir2, Chp2, Clr1) due to mis-annotations of the gene deletions or contaminations, as confirmed by genomic PCR and barcode sequencing (Suppl. Table S8; see Methods).

We also identified 189 gene deletions that caused a significant gain in silencing by applying reciprocal selection criteria (effect-size threshold < -2 for *MAT, SUBTEL*, and *TEL* and < -3 for *CEN*). The largest number of factors (116 genes) affected silencing at *CEN*, whereas fewer factors were found to regulate *MAT* (50) and *TEL* (54). Notably, only a few anti-silencing factors (4) were identified for *SUBTEL* (Figure 1b). Among factors counteracting silencing, we found components of the RNA polymerase-associated Paf1 complex (Paf1, Leo1), the histone H2B ubiquitin ligase complex HULC (Brl2, Shf1), and the RNA export factor Mlo3 (Suppl. Figure S4c), in agreement with previous reports (Flury et al., 2017; Kowalik et al., 2015; Reyes-Turcu et al., 2011; Sadeghi et al., 2015; Verrier et al., 2015; Zofall and Grewal, 2007). Interestingly, lack of the putative H3K9 demethylase Epe1, which prevents heterochromatin spreading beyond its boundaries (Ayoub et al., 2003; Zofall and Grewal, 2006), did not significantly affect silencing of any of the reporters in our study. This implies that the absence of Epe1 does not further increase *ura4+* silencing inserted at those heterochromatin loci, suggesting that intrinsic mechanisms that control Epe1 distribution on chromatin (e.g., Epe1 degradation) are sufficient to prevent its accumulation within heterochromatin in wild-type cells. In summary, our quantitative and systematic genome-wide screening approach retrieved a large number of factors that positively (180) and negatively (189) control silencing to different extents at constitutive heterochromatin domains.

### A range of factors regulate domain-specific heterochromatin silencing

Beyond the variability in the number of hits identified, we uncovered factors required to silence individual heterochromatin domains. Among the 180 mutants detected, only a small fraction (10 out of 180) affected silencing across all heterochromatin domains, while 26 and 36 impaired three or two regions, respectively (Suppl. Figure S5a, b). In contrast, 108 mutants, more than half, primarily affected a single heterochromatin domain (16 at *CEN*, 67 at *MAT*, 17 at *SUBTEL*, and 8 at *TEL*). Only *SUBTEL* and *TEL* exhibited a considerable degree of overlap. Remarkably, when comparing previous genome-wide screens conducted for individual heterochromatin loci (Bayne et al., 2014; Folco et al., 2019; Jahn et al., 2018; Kallgren et al., 2014; Kawakami et al., 2023; Wang et al., 2014b), we noticed similar patterns of limited overlap between hits for different heterochromatin domains (Supp. Figure S5b-d). However, the number of candidates undisclosed by our study exceeded those of previous studies at the individual scale and even when combined (see Discussion).

To further analyze domain-specific heterochromatin regulation, we performed k-means clustering and identified 7 distinct clusters, each showing a specific phenotypic profile (Figure 1c; Suppl. Figure S6a; Table 1; Suppl. Table S9). Cluster I displayed silencing defects throughout all heterochromatin regions and contained components mediating H3K9 methylation and spreading (CLRC and Swi6^HP1^). Cluster II exhibits similar defects yet weaker phenotypes for *TEL*. This cluster comprised additional components of the heterochromatin core machinery, including SHREC and the ubiquitin ligase Cul4-Ddb1^Cdt2^ mediating Epe1 degradation within heterochromatin. We also found several factors implicated in DNA replication, consistent with previous reports (Suppl. Table S10) (Jahn et al., 2018; Kawakami et al., 2023). Interestingly, while factors promoting RNAi are generally absent in Clusters I and II, the *arb2Δ* mutant differed significantly from other RNAi mutants, displaying a broad silencing defect that was also confirmed by RT-qPCR analysis (Suppl. Figure S7a; see Discussion).

**Table 1:**
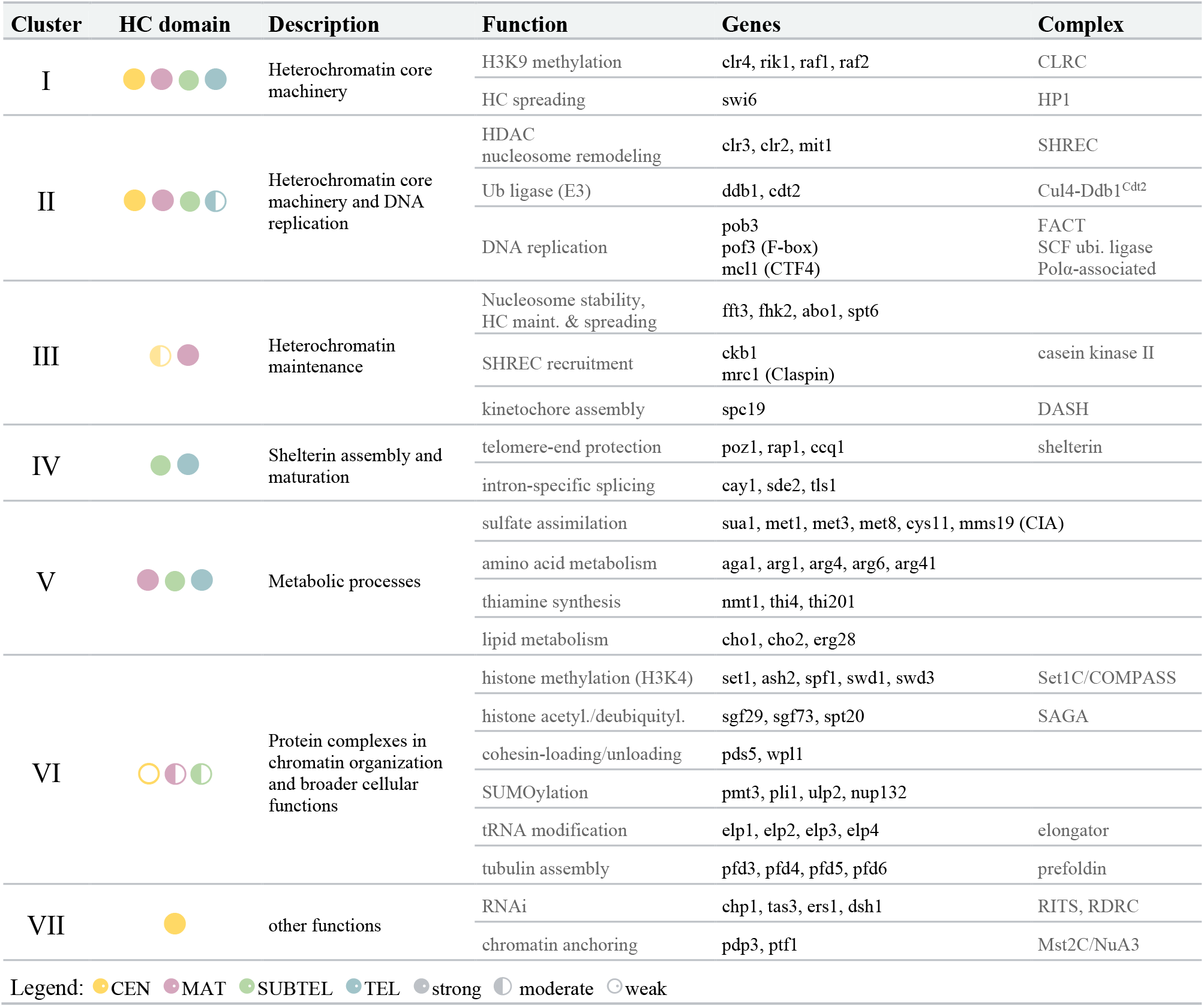
Clusters and representative mutants with decreased silencing.

Cluster III (15 mutants) exhibited a profound defect in *MAT* silencing, while displaying only subtle defects at other regions. Several factors present in this group have been linked to nucleosome stability and heterochromatin maintenance (e.g., the SMARCAD family nucleosome remodeler Fft3 (Taneja et al., 2017)) as well as SHREC recruitment (HP1 phosphorylation by casein kinase II (Shimada et al., 2009)). For others, the underlying mechanisms of silencing remain unclear. Notably, the loss of Spc19, a member of the DASH complex involved in kinetochore assembly, differed markedly from the moderate (or absent) phenotypes observed in other complex members, as confirmed by RT-qPCR (Suppl. Figure S7b). This difference suggests that Spc19 has a unique function independent of its role in DASH. In contrast, Cluster IV specifically affected *SUBTEL* and *TEL*. This group encompasses members of the telomere-protecting shelterin complex, including Ccq1, Rap1, and Poz1. We also found factors pivotal for the splicing of shelterin components. Notably, silencing in these mutants remained largely unperturbed at *CEN* and *MAT* (Suppl. Figure S7c), corroborating previous findings demonstrating their intron-specific roles in splicing the shelterin components Rap1 and Poz1 (Anil et al., 2022; Thakran et al., 2018; Wang et al., 2014b). Cluster V was similarly deficient in *SUBTEL* and *TEL* silencing but also displayed defects at *MAT*. Intriguingly, this relatively large group comprised many factors involved in various metabolic pathways (discussed further below).

Clusters VI and VII displayed less severe silencing defects (Suppl. Figure S6a). Cluster VI exhibits defects predominantly at *MAT* and *SUBTEL*. A distinctive feature was the presence of multiple protein complexes involved in chromatin regulation and broader cellular functions, such as Set1C/COMPASS (histone methylation), SAGA (histone acetylation and deubiquitylation), Elongator (tRNA modification), Prefoldin (tubulin assembly), and several factors involved in SUMOylation. While phenotypes in these mutants were subtle, they were highly reproducible and coherent among the complex members. Cluster VII exhibits primarily defects at *CEN* but weaker phenotypes at other regions. Prominent members of this cluster included components of the RITS and RDRC complexes, linked to RNAi, and the nuclear membrane protein Dsh1, which tethers these complexes to the nuclear periphery (Kawakami et al., 2012). This group also includes two components (Pdp3, Ptf1) of the Mst2C^NuA3^ histone acetyltransferase complex that anchor the complex to H3K36me3-marked euchromatin and prevent its encroachment into heterochromatin (Flury et al., 2017; Georgescu et al., 2020). Next, we aimed to uncover common characteristics among mutants that enhance silencing. These mutants showed less pronounced effects compared to those that decreased silencing. They further tended to manifest at the *CEN* reporter, consistent with the leaky expression of *ura4*^*+*^ at the *imr1L* locus and its strong repression at other heterochromatic regions in wild-type cells. To categorize the mutant phenotypes, we used k-means clustering, which resulted in 7 distinct groups (Table 2; Suppl. Figure S6b; Suppl. Table S9). The anti-silencing (AS)-Cluster I stood apart as displaying global defects with enhanced silencing at *CEN, MAT*, and partially at *TEL*. This group encompasses components of nucleosome remodeler Ino80C and factors involved in transcriptional elongation, pre-mRNA 3’-end processing, and mRNA export, in agreement with previous findings demonstrating that these processes counteract heterochromatin silencing (Flury et al., 2017; Kowalik et al., 2015; Reyes-Turcu et al., 2011; Sadeghi et al., 2015; Verrier et al., 2015; Yu et al., 2018; Zofall and Grewal, 2007). AS-Cluster II showed enhanced silencing predominantly at *CEN* and comprised the bromodomain protein Bdf2 and H2A.Z-specific histone chaperone SwrC, both preferentially associated with euchromatin (Iglesias et al., 2020). SwrC and Ino80 were further shown to prevent heterochromatin spreading (Greenstein et al., 2022). In contrast, AS-Cluster III displayed enhanced silencing primarily at *MAT* and comprised the H2B-specific ubiquitin ligase HULC and factors linked to nonsense-mediated mRNA decay. AS-Cluster IV exhibited a similar but weaker pattern and included several factors involved in autophagy.

**Table 2:**
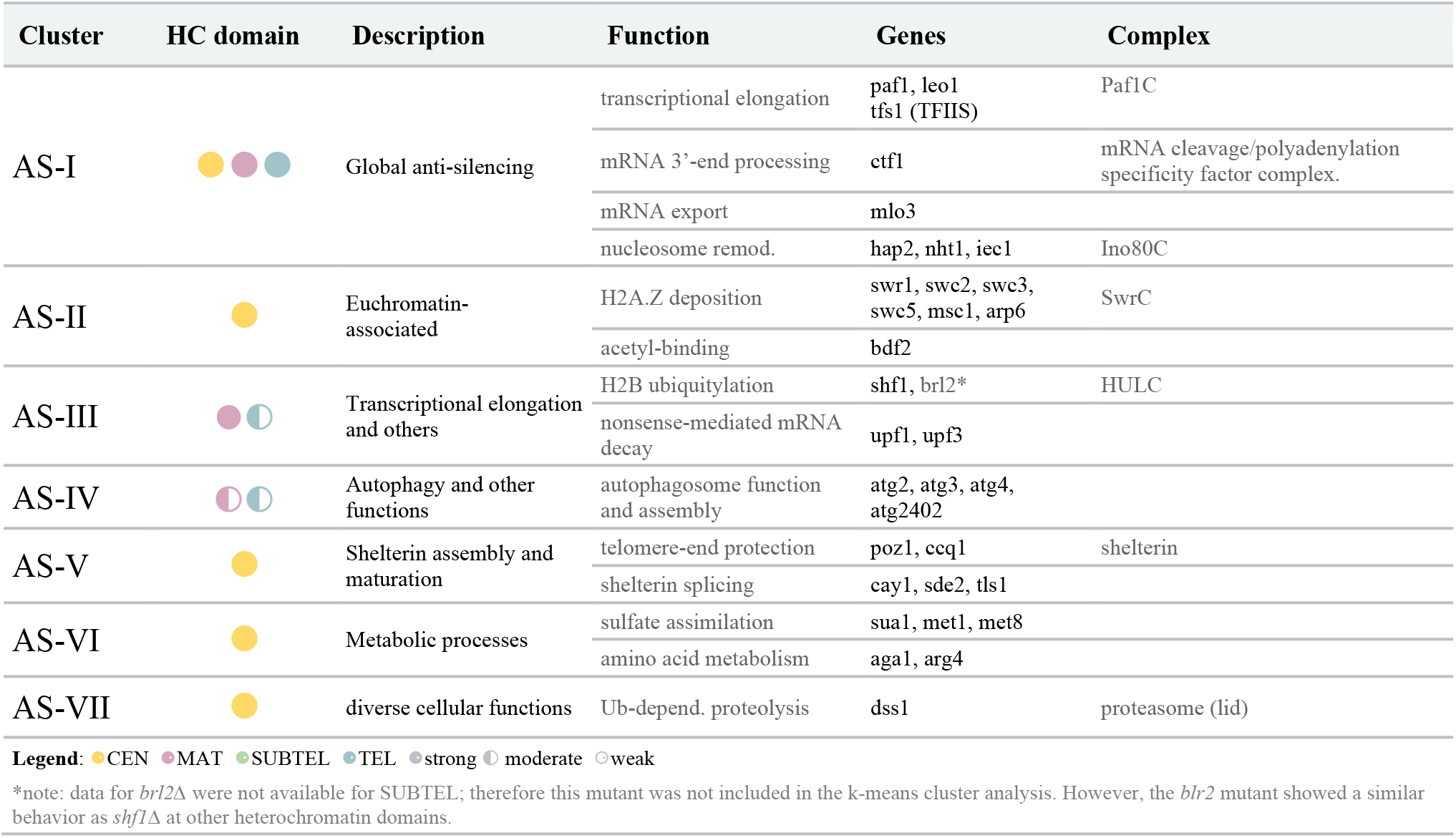
Clusters and representative mutants with enhanced silencing.

Intriguingly, several mutants with enhanced silencing at *CEN* displayed the opposite phenotype at *SUBTEL* and *TEL* (i.e., decreased silencing), and were therefore identified by both strategies. This was particularly evident for genes controlling shelterin composition and assembly (Silencing Cluster III, AS-Cluster V; Suppl. Figure S6a, b). These factors do not generally antagonize heterochromatin but rather have a context-specific role, consistent with previous reports (Barrales et al., 2016; Tadeo et al., 2013). Interestingly, we observed a similar trend for several mutants affecting metabolic pathways (Silencing Cluster V, AS-Cluster VI).

In conclusion, beyond identifying novel factors, the sensitivity of our quantitative screening approach and the generation of phenotypic profiles across different chromatin contexts let us assign factors to distinct pathways with heterochromatin region-specific functions.

### Phenotypic profiles reflect the submodular architecture of chromatin organization complexes

We noted strikingly similar phenotypic profiles among several components of physical protein complexes, such as CLRC, SHREC, and RITS (Figure 1c, d). This observation prompted us to systematically explore whether genetic perturbations in complexes associated with specific chromatin functions typically exhibit distinct phenotypic profiles. To this end, we calculated pairwise similarities between phenotypic profiles among 70 genes linked to 12 established chromatin organization complexes (Suppl. Table S11). Notably, genes encoding subunits of the same complex displayed higher similarities compared to those of different complexes (P<10^−4^ from permutation test, see Methods; Figure 2a). Specifically, 24.1% of within-complex pairs show a Pearson correlation coefficient of at least 0.9, whereas only 7.2% of between-complex pairs reached this level of correlation. This high level of correlation was particularly seen for members of the CLRC, SHREC, RITS, and shelterin complexes. However, we noticed that several larger protein complexes, such as Set1C/COMPASS, SAGA, and Mst2C^NuA3^, displayed greater diversity, with two or more gene correlation clusters within the same complex (Figure 2b). Focusing on these chromatin complexes, we investigated whether the mutant profiles align with the modular architecture of these complexes.

**Figure 2.**
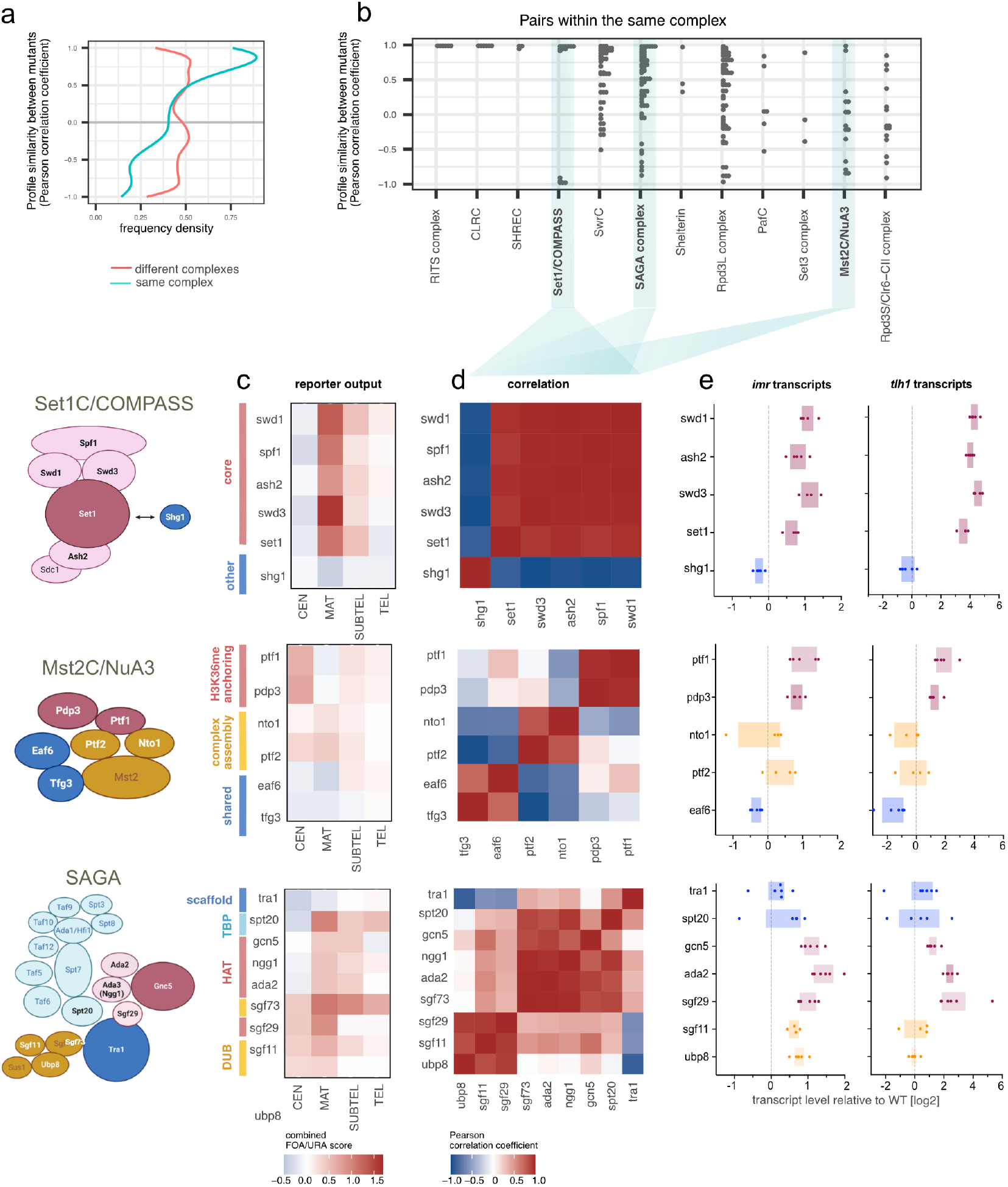
Similarities between phenotypic profiles reflect the composition of protein complexes involved in chromatin organization. a) Metablot showing frequency density of Pearson correlation coefficients for the pairwise correlation of mutant profiles of 70 genes belonging to 12 complexes involved in chromatin organization (for details, see Suppl. Table S11 and text). Green and red lines represent gene-pairs within the same or different complexes, respectively. b) Detailed view of pairwise correlations restricted to genes within identical complexes. Pearson correlation coefficients in (a) and (b) were calculated using average values of the combined FOA/URA score from 3-8 biological replicates. c) Heatmaps showing the median log2-transformed values of the combined FOA/URA scores from 3-8 biological replicates (*CEN*: 8; *MAT*: 7; *SUBTEL*: 3; *TEL*: 6; each with two technical replicates) focusing on three complexes: Set1C/COMPASS (top), Mst2C^NuA3^ (middle) and SAGA complexes (bottom). Accompanying diagrams (left panels) depict the complex structures based on existing crystallographic data or functional genetics studies. d) Correlation matrices highlighting Pearson correlation coefficients based on combined FOA/URA scores among subunits within each complex. e) Expression analysis of endogenous transcripts at two heterochromatic loci (*imr1L* and *tlh1*) analyzed by RT-qPCR in selected mutants. Transcript levels are normalized against *act1* levels and presented as box plots using log2-transformed values relative to the wild-type (WT) median across biological replicates (*n* = 4-5). Colors correspond to the different protein complex modules.

Based on the reporter growth data (Figure 2c; Suppl. Figure S8), correlation matrices were generated for each complex (Figure 2d). For the Set1/COMPASS histone H3K4 methyltransferase complex (Qu et al., 2018), the loss of its core subunits (Set1, Spf1, Swd1, Swd3, Ash2) resulted in highly similar phenotypic profiles (Figure 2d, top). Notably, these profiles markedly differed from that of Shg1, a protein that is dispensable for H3K4 methylation and only peripherally associated with Set1C (Mersman et al., 2012). A similar pattern was observed when examining endogenous heterochromatic transcripts by RT-qPCR, revealing an increase in core subunit mutants while no change was seen in cells lacking Shg1 (Figure 2e, top). This indicates that Shg1 is dispensable for heterochromatin regulation. Similarly, distinct phenotypic profiles were identified among subunits of the Mst2C/NuA3 acetyltransferase complex, responsible for modifying histone H3K14 and other chromatin-associated proteins (Figure 2d, middle) (Flury et al., 2017; Georgescu et al., 2020; Wang et al., 2012). Although the structure of this HAT complex remains elusive, correlating these phenotypic profiles revealed specific clusters (Figure 2d middle), which were also reflected by changes in heterochromatic transcript levels (Figure 2e, middle). Notably, these clusters corresponded with the roles of individual subunits in various functions: complex assembly (Nto1, Ptf1), anchoring of Mst2C to H3K36me3-marked chromatin (Pdp3, Ptf1), and shared activities with other chromatin complexes (Eaf, Tfg3), suggesting a modular architecture for Mst2C. Finally, when examining the multi-functional transcriptional co-activator complex SAGA (Qu et al., 2018; Wang et al., 2020), phenotypic profiles of the individual also segregated into the functional modules, including HAT (histone acetylation), DUB (histone deubiquitylation), and the TF-binding module (Figure 2d,e, bottom). However, some noticeable deviations were also observed. The phenotypic profile of *sgf73*Δ differed significantly from other DUB subunits, causing stronger silencing defects. This finding is consistent with the additional role of Sgf73 in RITS assembly independent of its function inside the SAGA complex (Deng et al., 2015). Furthermore, we found that the phenotypic profile of Sgf29, which is part of the HAT module, correlated with components in the DUB module, suggesting additional roles of Sgf29 within SAGA.

In conclusion, our reporter-based phenotypic profiles not only offer crucial information about the roles of chromatin complexes but also provide insights into their submodular architecture and the functions of these modules in relation to silencing.

### Metabolic pathway genes regulate subtelomeric and telomeric silencing

As mentioned earlier, mutants in Cluster V exhibited a distinct profile with reduced silencing defects at *MAT, SUBTEL*, and *TEL*, while silencing at *CEN* was enhanced (Figure 1c; Suppl. Figure S6a, b). To obtain insights into the roles of these genes, we conducted a GO term analysis using the AnGeLi web-based tool (Bitton et al., 2015). The analysis revealed significant enrichments in various, partially overlapping metabolic processes, including amino acid metabolic processes (GO: 0006520), arginine biosynthetic processes (GO:0006526), sulfur compound metabolic processes (GO: 0006790), and thiamine metabolic processes (GO:0006772) (Suppl. Figure S9a; Suppl. Table S12).

Examining these genes and their interactions using the STRING database (https://string-db.org/; (Szklarczyk et al., 2023), we identified several genes (*sua1, met1, met3, met6, met8, cys11, mms19*) involved in distinct steps of sulfur assimilation and the biosynthesis of homocysteine, methionine, cysteine, and S-adenosyl-methionine (SAM) (Figure 3a, b; Suppl. Figure S9b-d). SAM serves as a universal methyl donor for methylation reactions and may be specifically limiting for H3K9 methylation in these mutants. Additionally, we identified two genes (*cho1, cho2*) encoding SAM-dependent methyltransferases involved in the last steps of phosphatidylcholine synthesis, suggesting the potential importance of membrane composition in the silencing of these heterochromatin domains (Figure 3a, b). Supporting this idea, cells lacking the endoplasmic reticulum (ER) protein Erg28, which tethers several enzymes involved in ergosterol synthesis to the ER membrane, exhibited a similar phenotypic profile (Figure 3a; Suppl. Figure S9d). Other genes in this cluster contribute to thiamine (*nmt1, thi4, thi201*) and arginine synthesis (*aga1, arg4, arg6, arg11, arg12, arg41*), or are generally involved in amino acid metabolism (*maa1, pha2, SPAC56E4*.*03*) (Suppl. Figure S9d). We also noted that three additional mutants associated with these pathways (*met10, cys2, thi2*) displayed similar phenotypic profiles (Figure 3a), although they were initially not identified as candidates due to our stringent selection criteria (as described above). Collectively, the shared distinct phenotypic profiles of these mutants suggest that these genes play specific roles in the silencing of subtelomeric heterochromatin and the mating type locus. This notion is further supported by the finding that genes from these metabolic pathways, while significantly enriched in Cluster V, are mostly absent in the other groups, as determined by Fisher’s exact test (Suppl. Figure S9e). Given the prominent roles of genes involved in SAM production and phospholipid biosynthesis, we focused on selected mutants and validated the silencing defects by examining endogenous transcripts from different heterochromatin regions (Figure 3c). Consistent with the growth-based reporter assays, RT-qPCR analysis revealed a moderate but reproducible increase (three to sixfold) in the subtelomeric *tlh1* transcript level in mutants deficient in sulfur assimilation (*cys11, met3, met8*) and lipid metabolism (*cho1, cho2, erg28*). Other heterochromatic transcripts (*cen-dg, mat-Mc*) were either less affected or unaffected, confirming that defects are chromatin context-dependent in these mutants.

**Figure 3.**
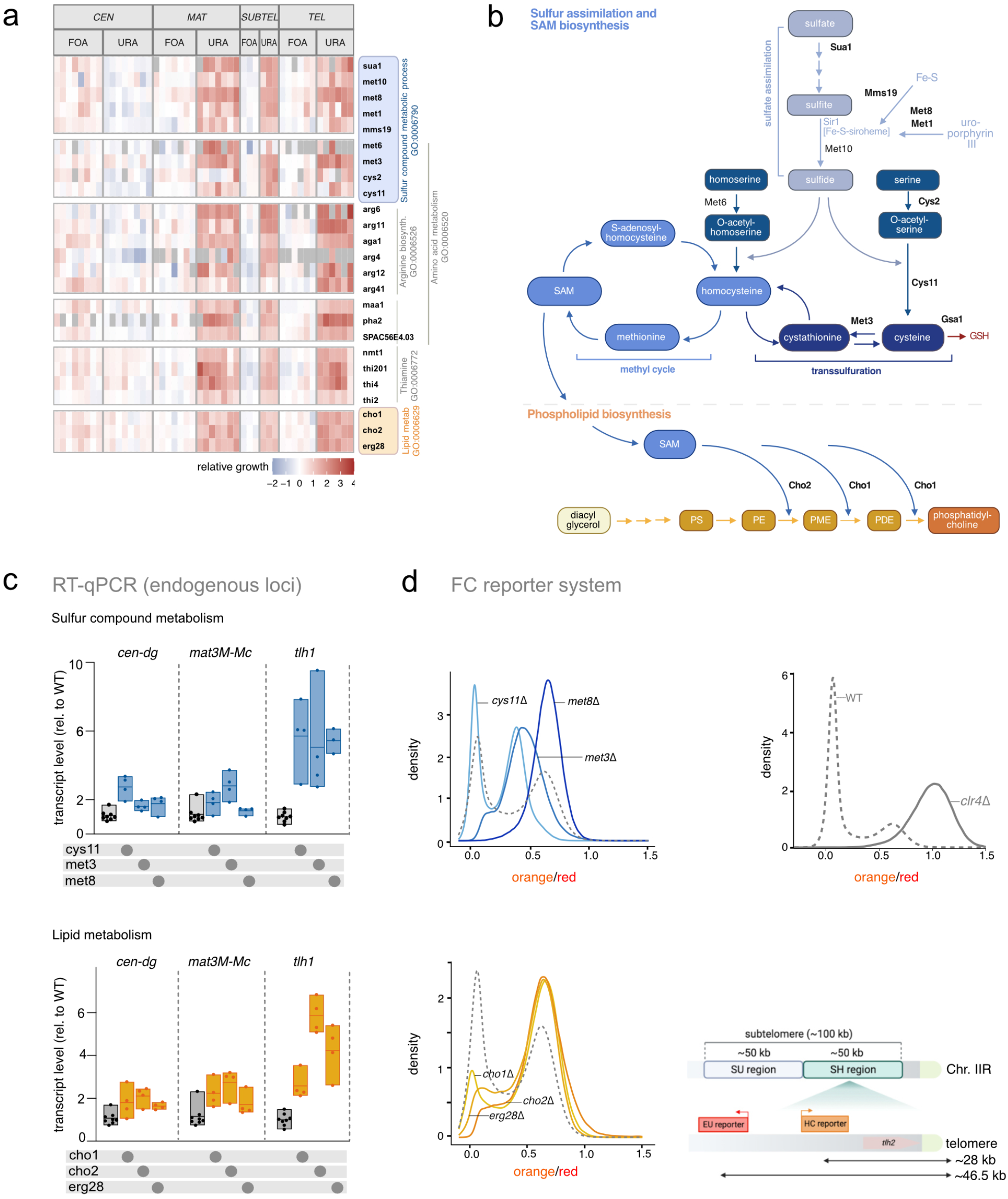
Metabolic pathway genes contribute to silencing at subtelomeres and mating type locus. **a)** Heatmap depicting relative growth values (+FOA, -URA) of mutants specifically impaired in silencing at *MAT, SUBTEL*, and *TEL*. Gene order reflects functional groups identified by Gene Ontology (GO) terms. **b)** Schematic representations of sulfate assimilation (top) and phosphatidylcholine synthesis (bottom) metabolic pathways. Bold/black protein names correspond to mutants identified under stringent selection criteria. **c**) Quantification of heterochromatin transcript levels at *cen-dg* (pericentromeres), *mat3M::ura4* (mating type locus) and *tlh1* (subtelomeres) by RT-qPCR. Transcript levels from 3-4 biological replicates, normalized against *act1* levels, are presented relative to the wild-type (WT) median (*n* = 8). Individual replicates are illustrated in a floating bar plot with the median indicated by a line. **d)** Density plots displaying flow cytometry analysis of fluorescent mKO2 reporter (‘orange’) expression at the single-cell level from a subtelomeric locus (HSS^Subtel^ reporter system). Flow cytometry experiments were conducted in mutants impaired in methionine and cysteine synthesis (top) and membrane lipid synthesis (controls: wild-type, dashed line; *clr4*Δ, solid line). The x-axis shows mKO2 reporter expression values (‘orange’) normalized against E2C expressed from a proximal euchromatic locus (noise filter, ‘red’). The y-axis represents the density of the cell population relative to the mean expression value in *clr4*Δ (ON state). The scheme illustrates the *HSS*^*Subtel*^ reporter system with ‘orange’ and ‘red’ inserted at ∼28 kb and ∼46 kb, respectively, at chromosome IIR.

To explore if bulk assays mask cell-to-cell differences in heterochromatin behavior in these mutants, we devised a fluorescent reporter system for measuring heterochromatin silencing in individual cells via flow cytometry (FC), as described previously (Greenstein et al., 2018). In this approach, we integrated a reporter gene encoding Kusabira Orange (‘orange’) 28 kb upstream of the telomeric repeats (Chr2R; Figure 3d). This specific locus exhibits lower levels of repression and is notably more sensitive to chromatin perturbations in comparison to other subtelomeric loci (A.Mazumder, B.A.-S., and S.B., unpublished results). To normalize signals from ‘orange’, we integrated an additional reporter, E2Crimson (‘red’), in nearby euchromatin. In a wild-type background, the reporter strain exhibited a bimodal behavior between a fully repressed (OFF) and an intermediately derepressed (ON) state, whereas it was completely de-repressed in cells lacking the methyltransferase Clr4. In the context of this reporter strain, mutants deficient in sulfate assimilation/homocysteine synthesis (*met3, met8*) displayed a pronounced shift towards the intermediate ON state, whereas cells deficient in cysteine synthesis (*cys11*) were less affected (Figure 3d, left top panel). Notably, lipid biosynthesis mutants (*cho1, cho2, erg28*) displayed an even stronger shift to the intermediate ON state (Figure 3d, left bottom panel). In addition to redistributing cells between ON and OFF states, some mutants additionally populate new intermediate ON states not found in wild type (*cys11* and *met3*). Therefore, together it stands to reason that homocysteine and phospholipid biosynthesis pathways are central to determining gene expression minima and maxima of bimodally distributed heterochromatin loci.

In summary, our phenotypic profiles uncover distinct and unanticipated roles for diverse metabolic pathways in the regulation of heterochromatin at subtelomeric domains. These regions appear to be more susceptible to cellular perturbations than other chromosomal regions.

### Dhm2 is a novel factor involved in silencing at constitutive and facultative heterochromatin

Among the mutants displaying broad heterochromatin defects in Cluster II (Figure 1c), we uncovered *dhm2* (deleterious haploid meiosis), encoding an uncharacterized protein of 11.25 kDa. Dhm2 is highly conserved among the *Schizosaccharomyces* clade (Figure 4a). Secondary structure prediction data from the AlphaFold consortium (Jumper *et al*, 2021) suggest that it consists of two alpha helices (Figure 4b). Dhm2 was previously identified in a sensitized genetic screening approach for mutants defective in heterochromatin maintenance at the mating type locus (Folco *et al*, 2019). However, its role in heterochromatin silencing remains unknown.

**Figure 4.**
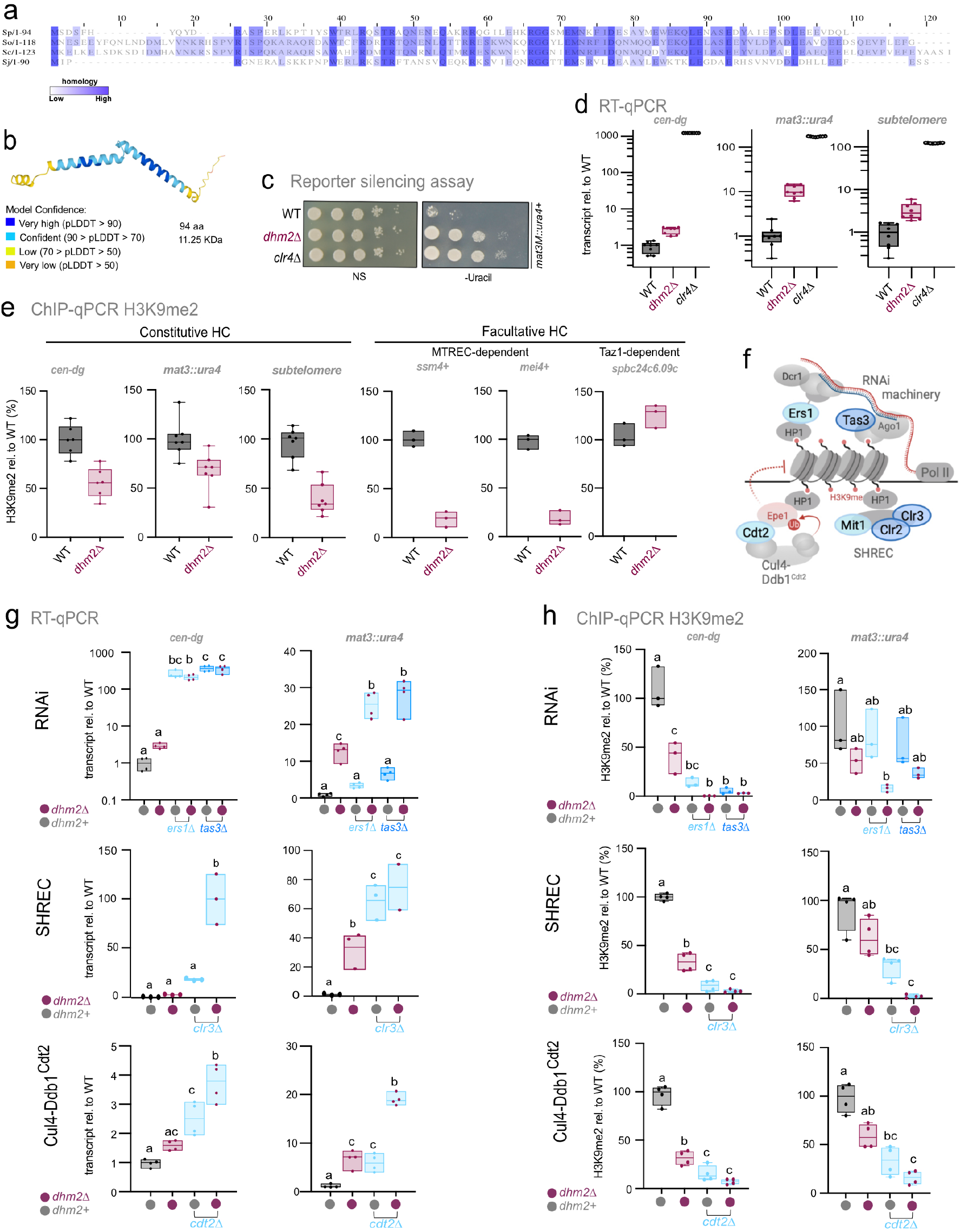
Dhm2 contributes to heterochromatin structure and silencing at constitutive and facultative heterochromatin. **a**) Multiple sequence alignment of the Dhm2 protein sequence and its homologs present in *S. osmophilus, S. cryophilus*, and *S. japonicus* using the T-coffee alignment tool. Residues are color-coded based on conservation levels (dark blue indicates high conservation). **b**) Predicted Dhm2 structure based on the AlphaFold2 model. **c**) Silencing assay using the *mat3M::ura4*^*+*^ reporter. Ten-fold dilutions of wild-type (WT) cells, *dhm2Δ*, and *clr4Δ* strains were plated on non-selective (N/S) and selective (lacking uracil) media. **d**) Quantification of heterochromatin transcript levels at *cen-dg* (pericentromeric repeats), *mat3M::ura4* (mating type locus), and *tlh1* (subtelomeric gene) by RT-qPCR. Transcript levels, normalized against *act1*, are presented relative to the WT median value (*n* = 8 independent biological replicates). **e**) ChIP-qPCR analysis of H3K9me2 levels at constitutive heterochromatin domains (*cen-dg, mat3M::ura4*, and *tlh1*^*+*^) and at facultative heterochromatin islands (*ssm4*^*+*^, *mei4*^*+*^, and *SPBC24C6*.*09c*). Input-normalized IP samples are normalized to the average of two euchromatic loci (*act1*^*+*^ and *tef3*^*+*^) and shown relative to the WT median value (n = 7 and 3 independent biological replicates, respectively). **f**) Schematic representation of heterochromatin pathways involving the RNAi machinery, SHREC, and Cul4-Ddb1^Cdt2^. **g**) Quantification of heterochromatic transcripts from *cen-dg* repeats and the *mat3::ura4*^*+*^ reporter gene by RT-qPCR. Transcript levels, normalized against *act1*, are presented relative to the WT median values (*n* = 3-4 independent biological replicates). **h**) ChIP-qPCR analysis of H3K9me2 enrichment at *cen-dg* repeats and the *mat3::ura4* reporter gene. Input-normalized IP samples, normalized to the average of two euchromatic loci (*act1*^*+*^ and *tef3*^*+*^), and are shown relative to the WT median value (*n* = 3-4 independent biological replicates). For **d, e, g and, h**, the individual replicates are displayed in box whisker or floating bar plots; the line depicts the median. For **g**, and **h**, statistical analysis was performed using one-way ANOVA tests, with letters denoting groups with significant differences as determined by Tukey’s *post hoc* tests at *P* < 0.05.

We therefore investigated the impact of Dhm2 on gene expression and the structural integrity of diverse heterochromatin domains. Using individual reporter growth assays and RT-qPCR analysis at the mating type locus, where the phenotype was most pronounced in data from our large-scale reporter assays, we confirmed the silencing defect in *dhm2*Δ cells, observing a 10-fold increase in *ura4+* gene expression (Figure 4c, d). Endogenous heterochromatic transcripts from pericentromeric repeats and subtelomeres were moderately increased (Figure 4d), corroborating the broad role of Dhm2 in heterochromatin silencing. To probe whether loss of Dhm2 affects heterochromatin structure, we performed chromatin immunoprecipitation with H3K9me2- and H3K9me3-specific antibodies. ChIP-qPCR revealed a partial H3K9me2 decrease at various constitutive heterochromatin domains, while H3K9me3 remained unaltered (Figure 4e; Suppl. Figure S10a). The absence of Dhm2 had a stronger effect at facultative heterochromatin, leading to a nearly complete loss of H3K9me2 at *ssm4*^*+*^ and *mei4*^*+*^, along with several other heterochromatin islands whose assembly requires the RNA processing and elimination factor MTREC (Figure 4e, Suppl. Figure S10b). In contrast, at other heterochromatin islands that are MTREC-independent but require Taz1 for assembly (e.g., *SPBC24c6*.*09*), H3K9me2 levels were unaffected or even increased (Figure 4e). Together, this implies a critical role of Dhm2 at specific facultative heterochromatin regions, whereas it appears to act redundantly with other pathways at constitutive heterochromatin domains.

To explore further a potential redundant role of Dhm2 in heterochromatin silencing, we introduced *dhm2*Δ into mutants lacking factors of known heterochromatin pathways (Figure 4f). When combining *dhm2*Δ with deficiency in RNAi (*ers1*Δ, *tas3*Δ), histone deacetylation (*clr3*Δ), or Epe1 degradation (*cdt2*Δ), we observed a strong synthetic defect in silencing (Figure 4g, Suppl. Figure S10c). In accordance with the aggravated silencing defects, H3K9me2 was virtually lost in these double mutants (Figure 4h, Suppl. Figure S10d). Together, these findings imply that Dhm2 contributes to heterochromatin silencing independently of these heterochromatin pathways.

### Dhm2 is required for heterochromatin maintenance

Perturbation in gene silencing due to the absence of Dhm2 could result from defects in either heterochromatin establishment, spreading, or maintenance. Initiation of heterochromatin establishment involves DNA-binding factors or RNAi (Grewal, 2023). To determine whether Dhm2 is required for RNAi-mediated silencing, we employed the previously described Rik1-λN/boxB reporter. Heterochromatin assembly is triggered through the recruitment of the RITS complex by Rik1, which requires an intact RNAi-machinery (Gerace et al., 2010). In this system, Rik1 is fused to the λN peptide recognizing boxB-binding elements sites integrated at the 3’-UTR of the *ura4* mRNA (ura4-5boxB; Figure 5a). As positive controls, we used mutants lacking Dcr1 or Mkt1, previously shown to be required for RNAi-dependent heterochromatin establishment (Taglini et al., 2020). In contrast to *dcr1*Δ and *mkt1*Δ, the loss of Dhm2 did not disrupt *ura4*^*+*^ silencing (Figure 5a). This result implies that Dhm2 is dispensable for RNAi-mediated post-transcriptional silencing, consistent with the above conclusion that Dhm2 acts redundantly with RNAi (Figure 4g, h).

**Figure 5.**
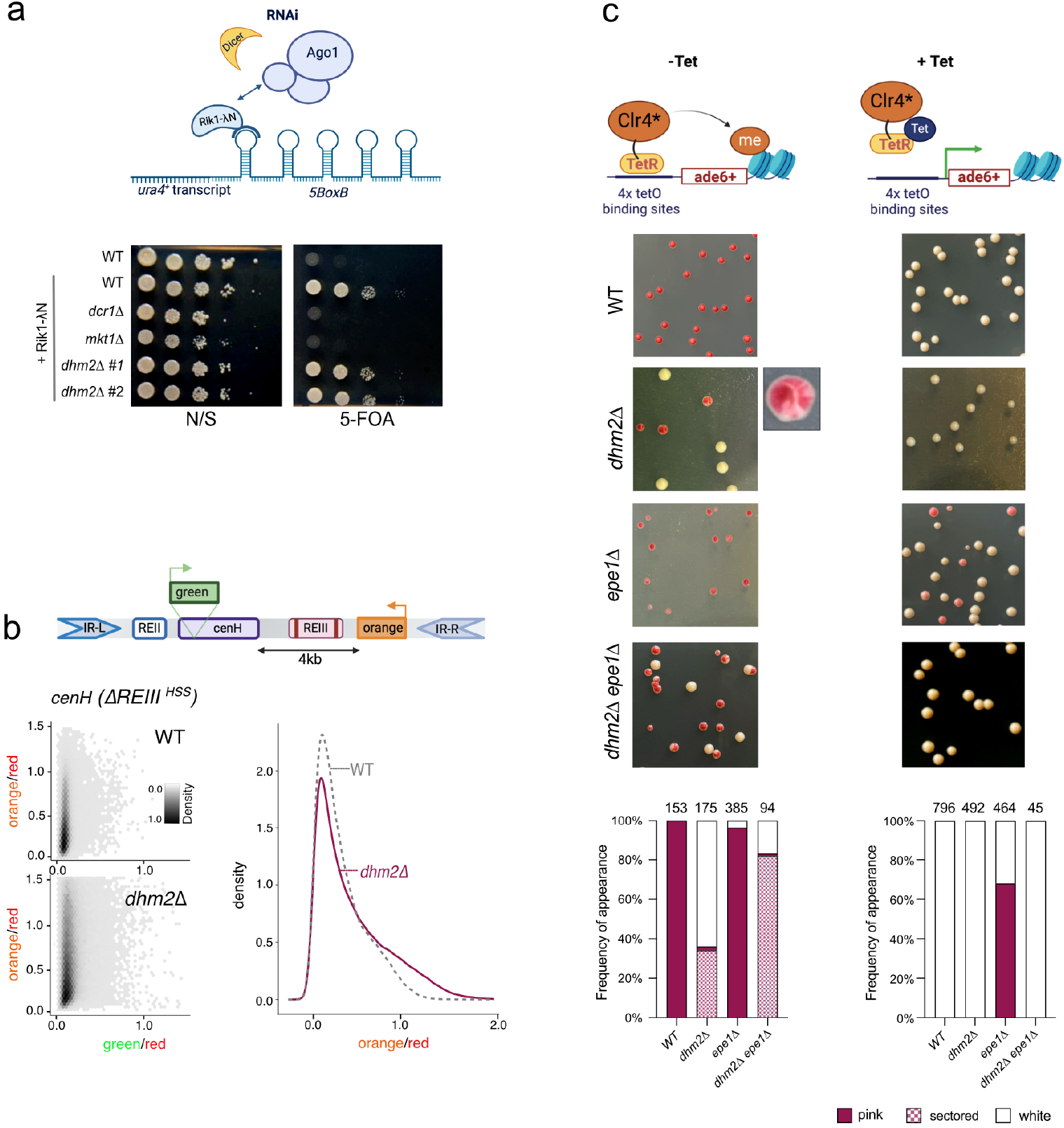
Dhm2 is required for heterochromatin maintenance. **a)** Monitoring of RNAi-dependent heterochromatin establishment. Top: Schematic of the Rik1 tethering system to nascent transcripts via boxB binding sites at the 3′UTR of *ura4*^*+*^. Bottom: Silencing assay using the *ura4*^*+*^*-5BoxB* reporter. Serial ten-fold dilutions of strains expressing Rik1-ΔN, including wild type (WT), positive controls (*dcr1Δ*, mkt1*Δ*), and two independent *dhm2Δ* strains, were plated on non-selective (N/S) and 5-FOA-containing media. The *rik1*^+^ strain expresses the non-fusion variant (negative control). **b)** Monitoring of heterochromatin spreading at the silent mating type locus. Top: Schematic of the Δ*REIII*^*HSS*^ (heterochromatin spreading sensor) system, with reporters inserted at *cenH* (‘green’; nucleation site), downstream of the REIII element (‘orange’; sensor site), and mutations in the Atf1/Pcr1 binding sites of REIII element (Δ*REIII*^*HSS*^) denoted by two vertical lines. An additional reporter gene (‘red’) is placed downstream of IR-R as a transcriptional noise filter (not shown). Bottom left: 2D hexbin plots display expression of ‘green’ and ‘orange’ reporters (normalized against ‘red’ expression), in WT and *dhm2*Δ mutant in the Δ*REIII*^*HSS*^ the reporter strain. Right: Density plot showing red-normalized ‘orange’ reporter expression cells filtered for ‘green’ off state. **c)** Monitoring of heterochromatin establishment and maintenance at an ectopic heterochromatin locus. Top: Schematic of the inducible TetR-Clr4* establishment system at the *ade6*^*+*^ locus. Middle: Representative images of colony color assay. WT, *dhm2*Δ, *epe1*Δ and *dhm2Δ epe1*Δ carrying *4xtetO-ade6*^*+*^ and expressing TetR-clr4*, were grown in the presence or absence of tetracycline (AHT). The graphs below show the percentage of red-pink colonies, sectored colonies, and white colonies. Absolute numbers of cells examined are indicated above the graph.

To further investigate the role of Dhm2 in RNAi-dependent and -independent heterochromatin establishment and spreading, we conducted assays at various heterochromatin loci. The MAT locus is a well-established region where heterochromatin is nucleated by the *cenH* element (homologous to the RNAi-nucleated pericentromeric *cen-dh* fragment) and the RNAi-independent *REIII* element. Both elements independently recruit the H3K9 methyltransferase Clr4, and once nucleated, heterochromatin spreads convergently from each element and is redundantly maintained by different pathways (Jia et al., 2004; Kim et al., 2004). However, mono-nucleated spreading can be studied by inactivating the *REIII* nucleation site so that spreading is initiated solely by the *cenH* element (Figure 5b). To examine nucleation and spreading, we employed the previously established heterochromatin spreading sensor (HSS) system, which uses three distinct fluorescent reporter genes inserted at a nucleation site (‘green’), a sensor site (‘orange’), and an unrelated locus outside heterochromatin for normalization (‘red’; see above) (Greenstein et al., 2018). This system can be used to measure both heterochromatin establishment at the nucleation site (OFF or ON state of ‘green’) and heterochromatin spreading by determining the ratio between the repressed state of ‘green’ (OFF) and ‘orange’ (ON or OFF). We employed the HSS system to study heterochromatin at different domains, including the mating type locus (*ΔREIII*^*HSS*^), subtelomeres (*SUB*^*HSS*^), and an ectopic locus (*EC*^*HSS*^) (see Methods).

We initially examined the ‘green’ reporter at the nucleation site in the ΔREIII^HSS^ strains to elucidate the role of Dhm2 in heterochromatin establishment. In *dhm2*Δ cells, we noted a minor population with increased green signal in comparison to the corresponding WT strain (Suppl. Figure S11a). This subtle effect was also evident in other domains, including two subtelomeric loci (∼11 and ∼37 kb upstream of the telomeric repeats) and an ectopic region, where heterochromatin is assembled via insertion of a pericentromeric *dh* element (Suppl. Figure S11b-d). To further investigate the influence of Dhm2 on heterochromatin spreading at the mating type locus, we analyzed the behavior of ‘orange’ in ΔREIII^HSS^ cells in which the green signal was OFF (indicating proper nucleation). Under this condition, a small subpopulation in ΔREIII^HSS^ cells gained the orange signal (Figure 5b). Together, these findings may suggest a minor contribution of Dhm2 to heterochromatin establishment and spreading. However, given the subtle nature of these changes, this may not be the primary cause of the global silencing defects we observe in the absence of Dhm2.

Therefore, we examined whether Dhm2 functions in heterochromatin maintenance using an inducible heterochromatin establishment assay that allows monitoring of heterochromatin maintenance once the initial trigger mediating establishment has been switched off (Audergon et al., 2015). In the absence of tetracycline (-AHT), the TetR-Clr4* fusion protein is recruited to *4xtetO* binding sequences, resulting in the silencing of the adjacent *ade6*^*+*^ reporter gene and, consequently, cells turning red on media containing low adenine. Conversely, upon the addition of tetracycline (+AHT), TetR-Clr4* dissociates from the *4xtetO* site, allowing *ade6* expression and formation of white colonies due to H3K9me turnover, promoted by the putative demethylase Epe1 (Audergon et al., 2015). We observed that, under establishment condition (-AHT), loss of Dhm2 resulted in increased white colony formation (more than 60%; Figure 5c). Many of the remaining red-pinkish *dhm2*Δ colonies exhibited a sectored phenotype, akin to mutants with defective clonal propagation of heterochromatin during cell division (Holla et al., 2020; Taneja et al., 2017). Notably, the appearance of white colonies, but not of the sectored phenotype, was partially suppressed in the *dhm2*Δ *epe1*Δ double mutant. Under the maintenance condition (+AHT), both WT and *dhm2Δ* cells displayed 100% white colonies (Figure 5c), consistent with previous reports that heterochromatin at this locus cannot be maintained in the presence of Epe1 that counteracts H3K9me (Audergon et al., 2015; Ragunathan et al., 2015). While the repressed state was partially retained in *epe1*Δ cells (65% red colonies), this was not observed in the *dhm2*Δ *epe1*Δ double mutant (100% white colonies), suggesting that Dhm2-dependent heterochromatin maintenance is independent of Epe1. Overall, these findings argue that Dhm2 plays a critical role in heterochromatin establishment and maintenance at this ectopic locus.

### Loss of Dhm2 results in replication stress

Previous studies have identified various DNA replication factors contributing to heterochromatin silencing (Jahn et al., 2018; Kawakami et al., 2023; Li et al., 2011), consistent with the notion that the propagation of histone modifications is crucial for epigenetic inheritance during cell division. Our study also identified several replication factors and confirmed their requirement for silencing various heterochromatin transcripts (Suppl. Figure S12a, b). Among these factors was the F-box protein Pof3^Dia2^, which ubiquitylates DNA polymerases and other replication factors (Maculins et al., 2015; Mamnun et al., 2006; Mimura et al., 2009; Roseaulin et al., 2013; Takayama et al., 2010).

Interestingly, while the *pof3*Δ mutant exhibited stronger silencing defects than *dhm2*Δ cells, the double mutant *dhm2Δ pof3Δ* displayed non-additive defects for pericentromeric transcripts, as determined by RT-qPCR, suggesting similar functions or shared pathways (Figure 6a). This functional relationship was also observed for the decrease of H3K9me2 levels in the single and double mutants (Figure 6b). In contrast, at other heterochromatin domains (mating type locus, subtelomeres), we observed additive or synthetic silencing defects implying that these factors act independently at these regions (Figure 6a). Attempts to generate viable double mutants of *dhm2* with other replication factors (Mcl1, Mrc1) were unsuccessful (Suppl. Figure S12c), implying that the combinatorial loss is synthetically lethal. This further supports the notion that Dhm2 is functionally linked to DNA replication.

**Figure 6.**
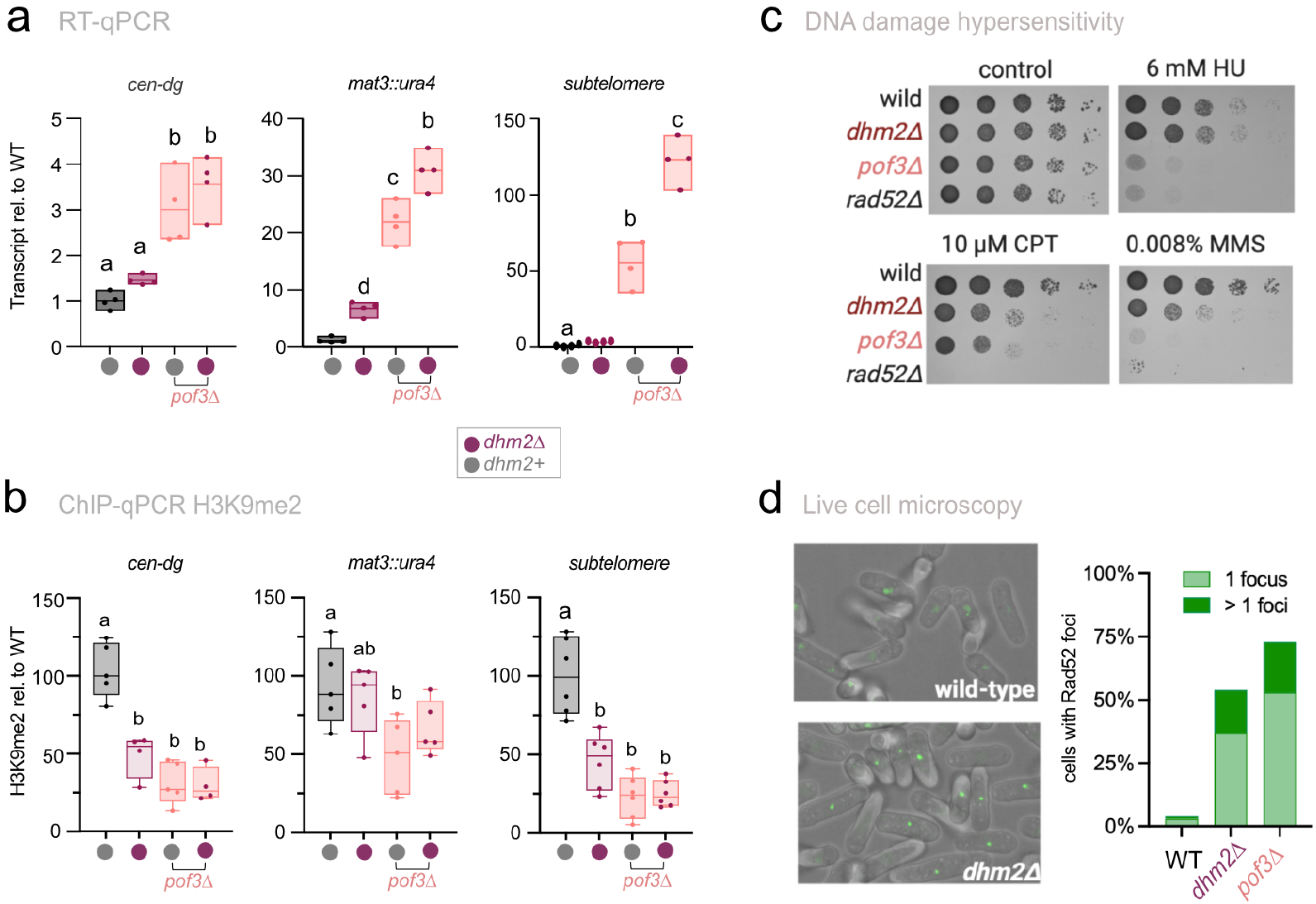
Loss of Dhm2 results in replication stress. **a**) RT-qPCR analysis of heterochromatic transcripts (*cen-dg, mat3M::ura4*, and *tlh1*) in the indicated strains. Transcript levels, normalized against *act1*, are presented relative to the WT median value (*n* = 4 independent biological replicates). **b**) ChIP-qPCR analysis of H3K9me2 levels at heterochromatin domains (*cen-dg, mat3::ura4*, and *tlh1*^*+*^) in the indicated strains. Input-normalized IP samples, normalized to the average of two euchromatic loci (*act1*^*+*^ and *tef3*^*+*^), are shown relative to the WT median value (*n* = 4-6 independent biological replicates). **c**) Sensitivity to DNA damage. Tenfold serial dilutions of the indicated strains were plated on YES medium supplemented with different DNA-damaging agents (HU, hydroxyurea; CPT, camptothecin; MMS, methyl methanesulfonate) and incubated for 3 days at 32°C. **d**) Accumulation of Rad52-GFP foci. Left panel: Representative images of WT and *dhm2*Δ cells expressing Rad52-GFP. Right panel: Percentage of Rad52-GFP foci formation in WT, *dhm2Δ*, and *pof3Δ* cells.

Many replication factors are sensitive towards genotoxic agents affecting various steps during replication fork progression. These include hydroxyurea (HU), which results in replication fork arrest through depletion of dNTP pools; camptothecin (CPT), which causes replication fork breakage by trapping topoisomerase on DNA; or methylmethane sulphonate (MMS), which blocks replication fork progression by DNA alkylation. Intriguingly, we found that *dhm2*Δ is sensitive toward CPT and MMS, causing similar growth defects to those seen for *pof3*Δ (Figure 6c). In contrast, *dhm2Δ* was insensitive toward thiabendazole (TBZ), an inhibitor of microtubule formation that affects mitotic progression in fission yeast (Suppl. Figure S12d). Rad52 (also known as Rad22 in *S. pombe*) plays a key role in DNA repair by homologous recombination (Ostermann et al., 1993). Upon induction of DNA double-strand breaks, Rad52 binds to single-stranded DNA, forming distinct nuclear foci (Lisby et al., 2001). Live cell imaging revealed that *dhm2*Δ displayed elevated levels of Rad52 foci in more than 50% of cells, similar to *pof3Δ* (∼70% of cells; Figure 6d), indicating that double-strand DNA breaks accumulate in the absence of Dhm2. Interestingly, Rad52 was also identified by our screens, and *rad52*Δ cells had a silencing defect at *MAT* (Cluster VI, Suppl. Figure S6a; Suppl. Figure S12e). Thus, in light of these findings, the defects in heterochromatin silencing and inheritance in *dhm2*Δ cells may result from defective DNA replication and/or DNA repair.

## Discussion

In this study, we present a systematic and quantitative growth-based reporter approach that unveils numerous factors influencing silencing at the major constitutive heterochromatin domains in *S. pombe*. The comprehensive nature of our investigation not only revealed many new factors but also corroborated the roles of genes previously associated with heterochromatin silencing, aligning our results with independent studies. Beyond the identification of factors, our quantitative approach and different reporter systems allowed us to allocate these factors to distinct functional pathways that show specificity for different heterochromatin domains. Among the new factors, we unveiled a potential role in heterochromatin maintenance and DNA replication or repair for Dhm2, providing a functional link between DNA replication and heterochromatin inheritance. In the following, we explore the implications of these findings.

### A comprehensive collection of factors implicated in heterochromatin silencing

From a pool of 100 genome-wide datasets, we identified 180 mutants that displayed a significant decrease in heterochromatin silencing, employing stringent selection criteria based on threshold and reproducibility. Many of the candidates we identified were further validated by directly examining heterochromatic transcripts, revealing a substantial consistency between growth-based reporter assays and transcript levels (Suppl. Figure S3, Suppl. Table S7). This collection of mutants represent a considerable increase in the number of candidates compared to previous studies employing the genome-wide gene deletion library from Bioneer. A key distinction in our approach was the use of multiple reporter strains to monitor silencing across different heterochromatin domains (*CEN, MAT, SUBTEL*, and *TEL*). In contrast, prior studies conducted single reporter screens (Bayne et al., 2014; Folco et al., 2019; Jahn et al., 2018; Kallgren et al., 2014; Kawakami et al., 2023; Wang et al., 2014b). Other critical factors contributing to the large number of candidates identified by our study include the use of quantitative measurements involving both technical replicates and multiple independent screening rounds, normalization of reporter assays, and the integration of two distinct readout methods for monitoring reporter activity (+FOA and -URA). This advanced approach provided our study with high robustness, reproducibility, and increased sensitivity in the context of reporter readouts.

In a few cases, we noticed some deviations between growth-based reporter assays and endogenous transcript levels. These may arise from differences in the experimental setup or sensitivity of assays, or could also be attributed to cellular heterogeneity. Using assays that enable single-cell detection, we could indeed segregate cellular subpopulations with different heterochromatin states (Figure 3).

Upon comparing published datasets with our study, we observed striking similarities with respect to specificity and overlapping functions of factors across heterochromatin domains. When consolidating all genes accumulated from previous studies, we found that approximately half of them (57 out of 120 genes) were also identified through our screens. It is noteworthy that the heterochromatin loci investigated in our study and previous reports were not entirely identical (Suppl. Figure S5).

While we investigated the pericentromeric *imr1L* repeats and a subtelomeric locus 7 kb upstream of telomeres, the other studies focused on the *cen-dg* repeats (Bayne et al., 2014; Kallgren et al., 2014) and the *SPAC212*.*07* gene 24 kb upstream of telomere (Kawakami et al., 2023), respectively. This experimental divergence could explain the differences in identified candidates. In contrast, the majority of genes identified for mating type silencing at *mat2P-ΔREII* and *mat3M-EcoRV* loci (Folco et al., 2019; Jahn et al., 2018) and telomeric repeats on the *Ch16* minichromosome (Wang et al., 2014b) were also discovered by our study, consistent with the identical or similar arrangement of the reporter genes (i.e., *mat3M-EcoRV*, telomeric repeats on *TEL2L*). Thus, independent datasets and variations in reporter systems increase the overall fidelity of our understanding of heterochromatin silencing.

### Specificity and requirement of regulators at different heterochromatin domains

A striking observation was that most candidates affected specific subsets of heterochromatin domains, while other regions remained unaffected (Figure 1, Suppl. Figure S5). This may reflect mechanistic differences in the establishment and maintenance of heterochromatin at those chromosomal regions. A notable example is RNAi, which is essential at pericentromeres but acts redundantly with additional pathways at other heterochromatin regions (Barrales et al., 2016; Hansen et al., 2005; Jia et al., 2004; Kanoh et al., 2005). Consistently, genes implicated in RNAi were exclusively identified through *CEN* reporter screens (Cluster VII; Figure 1, Suppl. Figure S6). An exception is the *arb2* mutant, lacking a component of ARC (Argonaute siRNA chaperone) involved in siRNA maturation (Buker *et al*, 2007). While this mutant exhibits a uniform silencing defect across all heterochromatin domains, this might be caused by additional misregulation of the juxtaposed *crb3* gene, which overlaps with the 3’-UTR of *arb2*^*+*^. Intriguingly, Crb3 is a member of the RNA-processing Rix1 complex (RIXC), and the combined loss of RIXC and RITS causes synthetic silencing defects (Holla et al., 2020; Shipkovenska et al., 2020). Another example of redundancy can be observed with the novel silencing factor Dhm2, whose role in silencing at constitutive heterochromatin is partially obscured by members of the heterochromatin core machinery (e.g., RNAi, SHREC). In contrast, the absence of Dhm2 results in the complete loss of heterochromatin at sites of facultative or ectopically induced heterochromatin, implying the lack of compensatory mechanisms in these regions (as discussed below). Heterochromatin domains may also vary in their specific needs in heterochromatin assembly. Spreading and maintenance of heterochromatin require mechanisms contributing to high nucleosome abundance and stability (Cutter DiPiazza et al., 2021; Greenstein et al., 2018; Taneja et al., 2017). Several factors promoting nucleosome stability (Fft3, Spt6, Abo1, and others) were enriched when selectively targeting silencing at the mating type locus (Cluster III and VI; Figure 1, Suppl. Figure S6). Notably, these factors were also identified by previous studies employing similar mating-type specific approaches (Folco et al., 2019; Jahn et al., 2018; Taneja et al., 2017). While some of these factors have been reported to affect additional heterochromatin regions (Gal et al., 2016; Jahn et al., 2018; Kiely et al., 2011), the pronounced sensitivity of the mating type locus suggests that this chromosomal region heavily relies on factors ensuring stable heterochromatin maintenance. A distinguished feature of this heterochromatin domain is the presence of multiple, well-defined nucleation sites, such as the *REII* and *REIII* elements, from which heterochromatin spreads into neighboring regions. In contrast, silencing at heterochromatin regions that primarily rely on RNAi, like the pericentromeric *dg* repeats, appears to undergo continuous heterochromatin establishment driven by siRNAs (Greenstein et al., 2018). Hence, these distinct mechanisms may also account for the requirement of different factors for heterochromatin maintenance.

Another chromatin context-specific observation was the identification of numerous factors linked to metabolic pathways, including methionine and membrane lipid biosynthesis, specifically affecting silencing at subtelomeres and the mating type locus (Figure 3; Suppl. Figure S9). While most of these factors were missed by the previous genome-wide studies, other research reported several links to methionine and SAM synthesis. Methylenetetrahydrofolate reductase Met11 plays a role in methionine regeneration through 5-MTHF generation and was noted to affect heterochromatin integrity (Lim et al., 2021). Additionally, SAM synthetase was among the SU(VAR) mutants displaying altered position effect of variegation in flies (Larsson et al., 1996) and was further shown to be crucial for silencing and perinuclear heterochromatin anchoring in worms (Towbin et al., 2012). Of note, in *S. pombe*, SAM synthetase is encoded by a single essential gene (Hayashi et al., 2018), explaining its absence in our study. Given SAM’s role in various methylation reactions, including H3K9me, methionine availability likely affects silencing in *S. pombe*, as observed in mammals (Mentch et al., 2015). SAM is also essential for the biosynthesis of membrane phospholipids. Supporting the idea of nuclear membrane composition impacting silencing, our study identified two methyltransferases (Cho2, Cho1^Opi3^) involved in phosphatidylcholine synthesis. Cho2 was previously also shown to physically interact with Lem2, another inner nuclear membrane protein contributing to heterochromatin silencing through multiple functions (Barrales et al., 2016; Hirano et al., 2023; Kinugasa et al., 2019; Tange et al., 2016). Thus, SAM deficiency appears to affect multiple pathways critical for silencing of subtelomeres (and to a lesser extent the mating type locus). Surprisingly, deficiencies in these pathways did not seem to impact pericentromeric heterochromatin (Figure 3). As different heterochromatin domains compete for shared pools of silencing factors (Barrales et al., 2016; Tadeo et al., 2013), this may suggest that subtelomeres are less favorable sites for heterochromatin assembly compared to other regions. Alternatively, the absence of defined boundary elements and the gradual decline of H3K9me-marked heterochromatin toward telomere-distal regions may cause subtelomeres to acquire dispersed heterochromatin structures, making them more vulnerable to fluctuations in the supply of factors required for their assembly, like SAM. In agreement with the chromatin source-sink hypothesis (Murphy and Berger, 2023), both scenarios are supported by the notion that subtelomeres can function as sinks for extra silencing factors, whereas the buffering capacity at pericentromeres is restricted by the presence of strict boundaries (Tadeo et al., 2013; Wang et al., 2014a).

### Unveiling additional functions of heterochromatin regulators

Our systematic approach comprehensively captured previously known architectural and functional submodules in protein complexes involved in chromatin organization (Figure 2). However, we also identified notable exceptions, suggesting additional roles for these factors. This was evident for complex members of SAGA (Sgf29, Sgf73), DASH (Spc19), or Mms19, a component of the cytosolic iron-sulfur assembly (CIA) machinery (Figure 2, Suppl. Figure S7). The DUB member Sgf73 was indeed shown to promote RITS assembly independently of its role within SAGA (Deng et al., 2015). Proteins from other complexes may share similar ‘moonlighting functions’ or play multiple roles in heterochromatin silencing. For instance, Mms19 has been linked to DNA metabolism and methionine biosynthesis (Gari et al., 2012; Kassube and Thomä, 2020; Stehling et al., 2012), but also associates with Cdc20 and Rik1-Raf2, factors linked to DNA replication and H3K9 methylation (Li et al., 2011). While these functions may not be mutually exclusive, the phenotypic profile of *mms19 Δ* resembles more closely those of mutants deficient in methionine biosynthesis (Figure 1, Cluster I vs. V). Moreover, Cdc20 possesses a 4Fe-4S cluster in its catalytic domain, suggesting that it might be a substrate of Mms19 which inserts Fe-S into apo-proteins (Jain et al., 2014). Therefore, further work is necessary to discern whether Mms19 functions directly (Rik1-Raf2 interaction) or indirectly (Fe-S cluster assembly) in heterochromatin maintenance.

Consistent with a previous proteomics study (Iglesias et al., 2020), our study identified two nucleoporins (Npp106, Nup132) and we confirmed their role in silencing heterochromatic transcripts (Suppl. Figure S6). While a broader involvement of nucleoporins in heterochromatin organization has been proposed, other inner nuclear membrane proteins present in Swi6^HP1^-purified heterochromatin did not appear to be crucial for silencing (Suppl. Table S3, Suppl. Figure S6) (Iglesias et al., 2020). Recently, Nup132 has been implicated in recruiting the SUMO protease Ulp1 to de-SUMOylate Lem2, a regulatory switch crucial for its role in silencing (Barrales et al., 2016; Strachan et al., 2023). Cells lacking Nup132 or Lem2 display silencing defects under rich growth conditions but not in minimal media, as used in our study (Martín Caballero et al., 2022; Strachan et al., 2023; Tange et al., 2016), explaining the absence of Lem2 in our current candidate list. The mechanism allowing cells to bypass Lem2 under certain conditions remains unclear, but this finding underscores the adaptability and dynamic regulation of heterochromatin pathways in diverse environmental conditions. Nevertheless, the retrieval of Nup132 in our study suggests additional functions in silencing. Thus, further exploration is needed to gain a comprehensive understanding of the role of nucleoporins in heterochromatin silencing.

### Role of Dhm2 and replication factors in epigenetic inheritance

Among mutants affecting all heterochromatin domains, we discovered Dhm2, a protein of unknown function previously identified in a genetic screen for MAT locus silencing defects (Folco et al., 2019). Given the lack of known protein motifs, the mechanism through which Dhm2 contributes to heterochromatin silencing remains unclear. We demonstrate that Dhm2 acts redundantly with various common silencing pathways at constitutive heterochromatin (Figure 4), providing a rationale for the moderate silencing defects observed in the single mutant. This may also explain why *dhm2*Δ had not been retrieved by most other studies.

Dhm2 may be involved in specific steps during heterochromatin assembly (i.e., nucleation, spreading, or maintenance). We found that Dhm2 is largely dispensable for RNAi- and shelterin-dependent heterochromatin establishment, consistent with acting redundantly rather than through these pathways. While we did not explore other possible RNA- or DNA-dependent establishment mechanisms, the broad involvement of Dhm2 at diverse heterochromatin domains makes a specific role in establishment less likely. We also observed only a modest effect on heterochromatin spreading when examining the mating-type locus (Zhang et al., 2008). However, Dhm2 had a substantial impact on heterochromatin maintenance at an ectopic locus where heterochromatin assembly can be induced independently of RNAi (Figure 5). Although silencing at this locus was compromised under both heterochromatin establishment and maintenance conditions, two critical observations suggest Dhm2 primarily contributes to the latter. First, under heterochromatin establishment conditions, red colonies (repressed reporter gene) often displayed a red-white sectoring phenotype, indicating that heterochromatin cannot be stably maintained without Dhm2. This variegating phenotype, shared by many mutants deficient in heterochromatin maintenance (Holla et al., 2020; Shipkovenska et al., 2020; Taneja et al., 2017), suggests that heterochromatin is not properly inherited during cell division. The appearance of white colonies may further imply a quick turnover of heterochromatin rather than a defect in the initial establishment. Second, although the absence of Epe1 allows silencing even under maintenance conditions (i.e., when de novo heterochromatin assembly is absent), maintenance in *epe1*Δ cells was completely lost by the additional lack of Dhm2, resulting in the exclusive appearance of white colonies. This finding not only implies that Dhm2 is critical for heterochromatin maintenance but also that it acts independently of Epe1.

The loss of heterochromatin maintenance in *dhm2Δ* may result from defects in epigenetic inheritance during DNA replication. In line with prior reports (Jahn et al., 2018; Kawakami et al., 2023; Li et al., 2011; Nathanailidou et al., 2024), we identified various replication factors, although they differed in the extent of silencing defects and specificity of the affected heterochromatin domains (Suppl. Figure S12). The phenotypic profiles (Figure 1) and the partially non-additive genetic interactions of Dhm2 and the F-box protein Pof3^Dia2^ (Figure 6) suggest that they act in similar pathways. As part of a ubiquitin ligase, Pof3^Dia2^ mediates the turnover of DNA polymerases and other replication factors (Maculins et al., 2015; Mamnun et al., 2006; Mimura et al., 2009; Roseaulin et al., 2013; Takayama et al., 2010). Furthermore, its absence causes hypersensitivity toward various DNA-damaging agents and accumulation of Rad52 foci indicating dsDNA breaks, a phenotype partially shared by *dhm2*Δ (Figure 6). Thus, it is plausible that Dhm2 is directly involved in Pof3related steps, although both may also have independent functions in different chromatin contexts, as indicated by the synergistic defects at subtelomeric heterochromatin. Additional work is needed to determine the role and interactions between Dhm2 and Pof3.

Dhm2 is highly conserved within the *Schizosaccharomyces* species, but no homologs have been found outside this clade. However, its critical function in maintaining heterochromatin structures in the absence of redundant pathways makes it likely that similar mechanisms exist in other eukaryotes. The predicted alpha-helical structure and the lack of known protein motifs suggest that it acts as an adaptor, facilitating the interactions of partner proteins. Given its small size, it may be possible that structures similar to Dhm2 are integral components of other polypeptides in higher eukaryotes. Future work focused on identifying Dhm2 interaction partners and gaining insights into its proteome will elucidate its role in heterochromatin inheritance, its potential link to DNA replication, and the broader conservation of those mechanisms.

### Concluding remarks

Our present study provides critical insights into the diverse roles of functional pathways and their specific contributions to heterochromatin domains, a previously underexplored aspect of heterochromatin biology. By employing a combination of multiple, quantitative, and sensitive reporter systems and phenotypic profiling, we have identified a plethora of heterochromatin regulators that often exhibit domain-specific silencing effects. This approach has also led to the discovery of numerous genes involved in metabolic pathways and elucidated the role of the newly identified regulator Dhm2 in heterochromatin maintenance. Furthermore, we have uncovered distinct functions within physical protein complexes, submodules, or even individual subunits, suggesting the existence of ‘moonlighting’ activities for some of these factors. This wealth of knowledge represents a significant contribution to the field, offering valuable insights for future studies.

Despite the significant findings of our study, we recognize certain limitations. Genetic screens employing single mutants can only identify factors that play essential roles in silencing. The absence of a pronounced phenotype in a single mutant at a particular heterochromatin domain does not preclude its potential involvement in silencing. Thus, factors exhibiting domain-specific behaviors may also fulfill additional roles at other heterochromatin domains, masked by the presence of redundant pathways. Additionally, certain factors and pathways may be critical for silencing only under specific conditions. Finally, our study focused exclusively on constitutive heterochromatin domains, while other genomic regions, such as facultative heterochromatin, were excluded. This could explain the absence of conserved proteins implicated in regulating heterochromatin in other systems, including human cells (McCarthy et al., 2021). Therefore, we recommend future research incorporating follow-up screens utilizing additional reporters and combinatorial approaches, such as E-MAP (epistasis mini array profiling) and varying growth conditions, to address these limitations.

## Materials and Methods

### Yeast strains and media

A modified version of the Bioneer *S. pombe* haploid deletion mutant library (version 3.0), in which non-essential genes were replaced with a *kanMX* cassette, was used for the reporter screens. In this collection encompassing 2988 mutants, several mtDNA-encoded genes and mutants with severe growth phenotypes had been removed (**Suppl. Table S1**). All other *S. pombe* strains used in this study are listed in **Suppl. Table S14**. Gene deletions and yeast strains expressing epitope-tagged proteins were generated by standard genome engineering procedures using transformation with PCR products and genomic integration via homologous recombination, as described earlier (Janke et al., 2004). Generated strains were validated by colony PCR. Reporter strains described in **Figure 3d** were generated by inserting three transcriptionally encoded fluorescent reporters into the subtelomere of chromosome arm IIR into the SD4 strain from Junko Kanoh’s laboratory (Tashiro et al., 2016) using a CRISPR/Cas9-based method (SpEDIT) (Torres-Garcia et al., 2020). For the reporters, the gene sequences were codon-optimized for *S. pombe* (Al-Sady et al., 2016) and the following constructs were used: Superfolder GFP (SF-GFP_s.p._, “green”) driven by the *ade6* promoter was integrated proximal to the *tlh2* gene approximately 11 kb downstream of the telomeric repeats; *ade6p*-driven Kusabira orange (mKO2_s.p._, “orange”) was integrated at either ∼28 kb or ∼37 kb from the telomeric repeats; *act1p*-driven E2Crimson (E2C_s.p._, “red”) was inserted at the nearest euchromatic region ∼46.5 kb (note: subtelomeric positions are corrected for the ∼ 4 kb sequence missing at the end of chromosome IIR in the annotated chromosome sequence on www.pombase.org). Strains shown in **Figure 5b** are derivatives from previously described reporter strains (Greenstein et al., 2018). For RT-qPCR and ChIP-qPCR experiments, cells were grown in rich medium (Yeast Extract Supplemented, YES) at 30°C. For genetic screens, cells were grown in synthetic medium (Edinburgh Minimal Medium, EMM). EMM medium supplemented with 5-FOA contained 1g/L 5′-fluoroorotic acid.

### Genome-wide screen for heterochromatin factors

Genome-wide screens were performed as described earlier (Barrales et al., 2016; Verrier et al., 2015) with some modifications. Briefly, a haploid deletion mutant library (Bioneer, version 3.0) was crossed with strains harboring the *ura4*^*+*^ reporter gene at different heterochromatic loci by using RoToR HDA colony pinning robot (Singer Instruments). To enable selection of the *ura4+* reporter under conditions of transcriptional repression, the reporter strains carried an additional hygromycin resistance marker (*hphMX6*) inserted into euchromatin next to the heterochromatin locus, which ensures robust genetic linkage during genetic crosses **(Suppl. Figure S1b)**. Following mating, cultures were incubated at 42ºC for 4 days to eliminate unmated haploid and non-sporulated diploid cells. Germination was induced by plating spores on YES media containing G418 and hygromycin B. For the readout, cells were transferred onto EMM, EMM lacking uracil (EMM-URA) and EMM supplemented with 5-fluoroorotic acid (EMM+FOA). Colonies were photographed and sizes were measured by the Gitter R package (https://omarwagih.github.io/gitter/). Relative growth was calculated by dividing colony sizes measured on selective medium (EMM-URA or EMM+FOA) by the respective sizes measured on non-selective medium (EMM). To compensate for plate-dependent variations, all values were normalized to the median values of the individual 384-plates. Log2 values were used for clustering and data visualization. For every reporter, we screened the haploid deletion collection multiple times independently (6-8 biological replicates, except subtelomeres n = 3; each biological replicate contained two technical replicates from the same genetic cross generated by duplication during the germination step).

### Confirmation of gene deletions by PCR or barcode sequencing

Prominent mutants isolated as hits in the screens were tested for the correct deletion. PCR analysis of genomic DNA was used to detect the proper junction of the integrated cassette (*kanMX*) or the absence of the deleted ORF using gene-specific primers. Alternatively, the presence of the strain-specific deletion cassette was confirmed by barcode sequencing. Following purification of genomic DNA and PCR amplification using 5’-GCAGTTTCATTTGATGCTCG-3’ and 5’-TTGCGTTGCGTAGGGGGG-3’ as forward and reverse primers respectively, the barcode sequence (‘downtag’) was analyzed by sequencing and compared with sequences present in the database (http://pombe.kaist.ac.kr/nbtsupp/ or in Han et al. 2010 (Han et al., 2010). Misannotated deletion strains (listed in **Suppl. Table S8**) were either removed from the analysis or renamed using the correct gene (those mutants are annotated with an asterisk). In some instances, this resulted in two copies of the same mutant in our collection (the original mutant and the corrected mutant, e.g., *swi6* and *swi6\**); in this case, we kept the explicitly confirmed mutant (i.e., *swi6*\*). Mutants that were contaminated by other mutants (e.g., *clr1*, a subunit of SHREC) or false positives containing a *URA3* harboring plasmid initially used by *Bioneer* to generate the haploid deletion library (e.g., *mst2*, a histone acetylase complex subunit) were excluded from the analysis.

### Threshold settings and statistical methods

To determine the number of silencing and anti-silencing genes from the screens, we applied the following steps. First, we calculated the mean of the relative growth value from the two technical replicates, which we considered as biological replicate value. Next, to make these values comparable across the four different reporter screens, we scaled the datasets by setting the standard deviation to one and centered them by setting the median of the screens to zero (**Suppl. Figure S2**). Then, we combined the growth values derived from the two readouts (EMM-URA and EMM+FOA, considering that silencing defects appear as growth inhibition on FOA medium (resulting in negative log_2_ values) and increased growth on medium lacking uracil (positive log_2_ values). Thus, for combining both readouts, we first multiplied the FOA score by (−1) before adding together the two types of scores. We refer to this value as *combined FOA/URA score*. Combining both readouts was necessary, because several strains exhibited strong effects by one readout but weak effects by the other. Next, one-sample Student’s t-tests were used to identify mutants whose scores were significantly different from 0 (median of the screens). To define significant hits, we employed a P-value < 0.05 and an effect-size threshold for the median combined score of biological replicates > 2.5 (*MAT, SUBTEL, TEL*) or > 3 (*CEN*; a higher threshold was chosen for *CEN* due to the leaky expression of the *ura4+* at the *imr1L* locus). The validity of these thresholds was assessed by determining the sensitivity in retrieving known heterochromatin factors (recall) and analyzing the precision of ‘hits’ by examining transcript levels of *ura4+* directly by RT-qPCR (positive predictive value). To this end, we compiled a list of *bona fide* heterochromatin factors based on available GO functional classification of *S. pombe* genes (GO categories: “heterochromatin” (GO:0000792), “chromosome, telomeric region” (GO:0000781), “heterochromatin boundary formation” (GO:0033696) and a phenotypic term “decreased chromatin silencing at subtelomere” (FYPO:0004604) (**Suppl. Table S4**). By applying these selection criteria (P-value, effect-size threshold), we retrieved recall values of 78%, 67%, 73%, and 69% for the *CEN, MAT, SUBTEL*, and *TEL* screens, respectively (**Suppl. Figure S2b; Suppl. Table S6**). Since several *bona fide* factors (e.g., RNAi factors) act redundantly at *MAT, SUBTEL*, and *TEL*, they were therefore not detected by these heterochromatin reporters, explaining values of less than 75%. For determining the precision, we analyzed the transcript levels of *ura4+* at *CEN, MAT* and *TEL* by RT-qPCR for a representative subset of mutants retrieved by the reporter growth-based selection criteria (**Suppl. Figure S3)**. We found an elevated expression level (i.e., greater than 1.5-fold change compared to WT) in 50-92% of the tested mutants (depending on the heterochromatic region; note that *SUBTEL* was not tested; **Suppl. Tables S5** and **S6**).

### Generation of cluster groups

To define heterochromatin domain-specific features of the silencing regulators, we clustered the phenotypic profiles by applying the following steps. First, k-means clustering was performed by using the function ‘kmeans’ of the ‘stats’ R package. The number of clusters was estimated by considering the biological functions of the genes. To assess the reproducibility of cluster assignment, multiple rounds of k-means clustering (n > 10) were performed. In rare cases where genes could be assigned to more than one cluster, cluster assignment was made based on frequency or behavior of related genes (e.g., subunits of protein complexes). Next, hierarchical clustering was performed for each k-means cluster by using the function ‘hclust’ of the ‘stats’ R package by calculating Euclidean distance (**Figure 1c-d, Figure S6**). Both for the k-means and the hierarchical clustering, median values of the biological replicates were used, and values of the *SUBTEL* and *TEL* screens were half-weighted. For visualization of the heatmaps, values of the biological replicates (mean values of the technical replicates) were used. 4 mutants (out of the 180 hits) were excluded from the k-means clustering due to missing values.

R script is available at https://github.com/zsarkadi/Muhammad-Sarkadi-et-al.

### RNA extraction and cDNA synthesis

RNA extraction and cDNA synthesis were performed as previously described (Barrales et al., 2016; Braun et al., 2011). In brief, 50 mL of yeast cells (OD_600_ = 0.4-0.8) were centrifuged at 4°C and cell pellets were frozen in liquid nitrogen. Upon thawing on ice, cells were resuspended in 1 ml TRIzol reagent. Following the addition of 250 μL zirconia/silica beads (BioSpec), cells were lysed by bead beating (Precellys 24, Bertin instruments) for 3 x 30 seconds with 5 minutes rest on ice, followed by centrifugation at 13,000 rpm for 15 minutes at 4 °C. Recovered supernatant was extracted twice with chloroform and centrifuged at 13000 rpm at 4 °C for 10 minutes. Following precipitation with isopropyl alcohol, the pellet was washed twice with 75% EtOH, air-dried and resuspended in 50 μL of RNase free dH2O. RNA concentration and quality were measured using a spectrophotometer (NanoDrop™, Thermo Scientific). Resuspended RNA was treated with DNaseI (Ambion) for 1 hour at 37°C. The reaction was stopped by adding 6 μL of inactivation reagent. For cDNA synthesis, 5 μg of DNase-treated RNA was converted into cDNA by reverse transcription using oligo(dT)_20_ primer (50 μM) and 0.25 μL of superscript IV (Invitrogen) according to the manufacturer’s instructions.

### Chromatin immunoprecipitation (ChIP)

ChIP-qPCR was performed as previously described (Braun et al., 2011; Georgescu et al., 2020), with some modifications. Yeast cultures (100 mL) were grown in YES media to mid-log phase (OD_600_ = 0.6-0.8) at 30ºC. Cells were cross-linked by adding formaldehyde to a final concentration of 1% for 20 minutes at room temperature (RT) by gentle shaking. Cross-linking was stopped by adding glycine to a final concentration of 150 mM for 10 minutes at RT. Cells were washed twice with 50 mL ice-cold PBS (phosphate-buffered saline) and resuspended in 1 mL lysis buffer A (50 mM HEPES/KOH, pH 7.5, 140 mM NaCl, 1 mM EDTA, 1% Triton X-100 (v/v), 0.1% NaDeoxycholate (w/v)), supplemented with Roche protease inhibitors. Cells were lysed by bead beating (Precellys 24, Bertin instruments) for 6 x 30 seconds with 5 minutes rest on ice. Genomic DNA was sheared by sonification (Q800R1 sonicator, QSonica) for 30 minutes (30-second on/off cycles, 90% amplitude) at 4 °C and cell debris was removed by centrifugation at 14000 rpm at 4 °C for 10 minutes. Soluble chromatin fractions were incubated with antibodies (anti-H3K9me2, Abcam ab1220; anti-H3K9me3, Active Motif, catalog no 39161) overnight at 4°C, followed by the addition of 25 μL of Dynabeads ProteinG (Life Technologies). Samples were washed 3x for 5 minutes at RT with lysis buffer A, 3x with high salt buffer (lysis buffer A containing 500 mM NaCl), and finally with 3x wash buffer (10mM Tris/HCl pH 8.0, 250 mM LiCl,1 mM EDTA, 0.5 mM NP-40 and 0.5% NaDeoxycholate (w/v)) and once with TE buffer (10mM Tris/HCl pH 7.5, 10 mM EDTA). DNA was de-crosslinked and eluted from the antibodies with ChIP Elution buffer (50 mM Tris HCl pH 7.5, 10 mM EDTA, 0.8% SDS) at 95°C for 15 minutes, followed by 65 °C for 4 hours. Eluted DNA was treated with Proteinase K at 55ºC for 30 min and then purified with a ChIP DNA Clean and Concentrator kit (Zymo Research) according to the manufacturer’s instructions.

### Quantitative gene expression (RT-qPCR) and chromatin association (ChIP-qPCR) analysis

RNA converted to cDNA and immunoprecipitated genomic DNA were quantified by real time PCR using PowerTrack™ SYBR Green Mastermix (Applied Biosystems™) and a QuantStudio 3 or QuantStudio 5 Real-Time PCR instrument (Applied Biosystems™). Primers used for qPCR are listed in **Suppl. Table S13**. Relative expression values for WT and mutants were calculated by normalizing transcript levels to euchromatin control *act1* and then dividing by the mean of all samples from the same experiment (group normalization), as previously described (Georgescu et al., 2020). When analyzing single and double mutants for epistatic interactions, statistical testing was performed using R. Multiple testing was performed using oneway ANOVA followed by a Tukey’s *post hoc* test at a 0.05 significance level.

### Flow cytometry

Flow cytometry analysis was performed according to a previously described protocol (Greenstein et al, 2018). *S. pombe* cells were grown to stationary phase in rich media (YES) and then diluted to a concentration of OD_600_ = 0.1 in YES, followed by incubation at 32 °C for 4-5 hours prior to flow cytometry analysis. The BD Fortessa X-50 instrument (UCSF, San Francisco), equipped with a high-throughput sampler (HTS) module, was employed for flow cytometry analysis. Sample sizes ranged from approximately 2,000 to 100,000 cells depending on the growth conditions of the respective strain. Compensation was performed using strains expressing no fluorescent proteins (XFP) and a single-color control XFP (SF-GFPsp, mKO2sp, or E2Csp). Compensated SF-GFP and mKO2 signals were normalized to E2C expressed from a euchromatic control locus in each single cell. Additionally, a maximum expression value for SF-GFP and mKO2 was set based on their expression in a heterochromatin-deficient control strain (*clr4Δ*), where reporters should be in an ON state. However, since the reporters are inserted at heterochromatin domains which are prone to recombination in *clr4Δ* strains, there is a risk of losing these reporters. To overcome this issue, color-negative cells were excluded by setting a minimum cutoff for SF-GFP and mKO2 based on a control strain expressing only E2C that mimics a “fully repressed” state for both reporters. “Max” expression values for SF-GFP and mKO2 were then calculated from these color-positive cells in the no-heterochromatin control strain. Subsequently, SF-GFP and mKO2 signals were scaled to the corresponding “max” values in our analysis strains, and the scaled, normalized signals were plotted in 2D hexbin or density plots for visualization.

### Live cell microscopy

Live-cell imaging was essentially performed as described (Barrales et al., 2016). In brief, cells were grown overnight in rich medium (YES) to the logarithmic phase (OD_600_ = 0.4-0.6). Before imaging, cells were attached to lectin (Sigma) coated glassbottom dishes containing a microwell (MatTek). Cells were imaged using a Zeiss AxioObserver Z1 confocal spinning disk microscope with an EMM-CCD camera (Photometrics, Evolve 512) through a Zeiss Alpha Plan/Apo ×100/1.46 oil DIC M27 objective lens. *Z*-stacks were obtained at focus intervals of 0.4 μm. FiJi/ImageJ software was used to count the number of foci in the yeast cells.

## Availability of data and materials

The datasets supporting the conclusion of this article are included within the article and as Supplementary Tables S1-S14 as individual spreadsheets in a MS-Excel file.

Additional data and R scripts are available on the GitHub repository: https://github.com/zsarkadi/Muhammad-Sarkadi-et-al.

## Acknowledgments

We thank members of the Braun lab, A. Ladurner and S. Hake for fruitful discussions during the study. We thank R. Allshire, E. Bayne, J. Kanoh, and M. Knop for strains and plasmids. We thank S. Lall (Life Science Editors) for editorial assistance and critical comments on the manuscript. Furthermore, we thank S. Fischer-Burkart, M. Bingel, and T. Uhlig for technical assistance. We acknowledge support by the German Research Foundation (DFG) to S.B. through the Heisenberg Programm (project ID 464293512), the collaborative research center CRC 1064 (project ID 213249687-SFB1064), and individual grants (401430508, 505087133). Further support was provided to S.B. by the European Union through the Network of Excellence EpiGeneSys (HEALTH-2010-257082) and an MSCA ITN Cell2Cell fellowship to A.Maz. and V.N.S.S. (project ID 860675). B.A.-S. was supported by a National Institutes of Health grant R35GM141888 and a grant 2113319 from the National Science Foundation.

## Contributions

Conceived and designed the study: B.A-S., B.P., R.R.B., and S.B. Collected the data: A.Muh., Z.S., A.Maz., M.C., V.N.S.S., and R.R.B. Contributed analysis tools: T.v.E. Performed the data analysis. A.Muh., Z.S., T.v.E., A. Maz., M.C., G.F., V.N.S.S., and S.B. Wrote the paper: A.Muh., Z.S., R.R.B., and S.B. with input from all authors.

## Competing interests

The authors declare that they have no competing interests.

## Supplementary Figures

**Figure S1.**
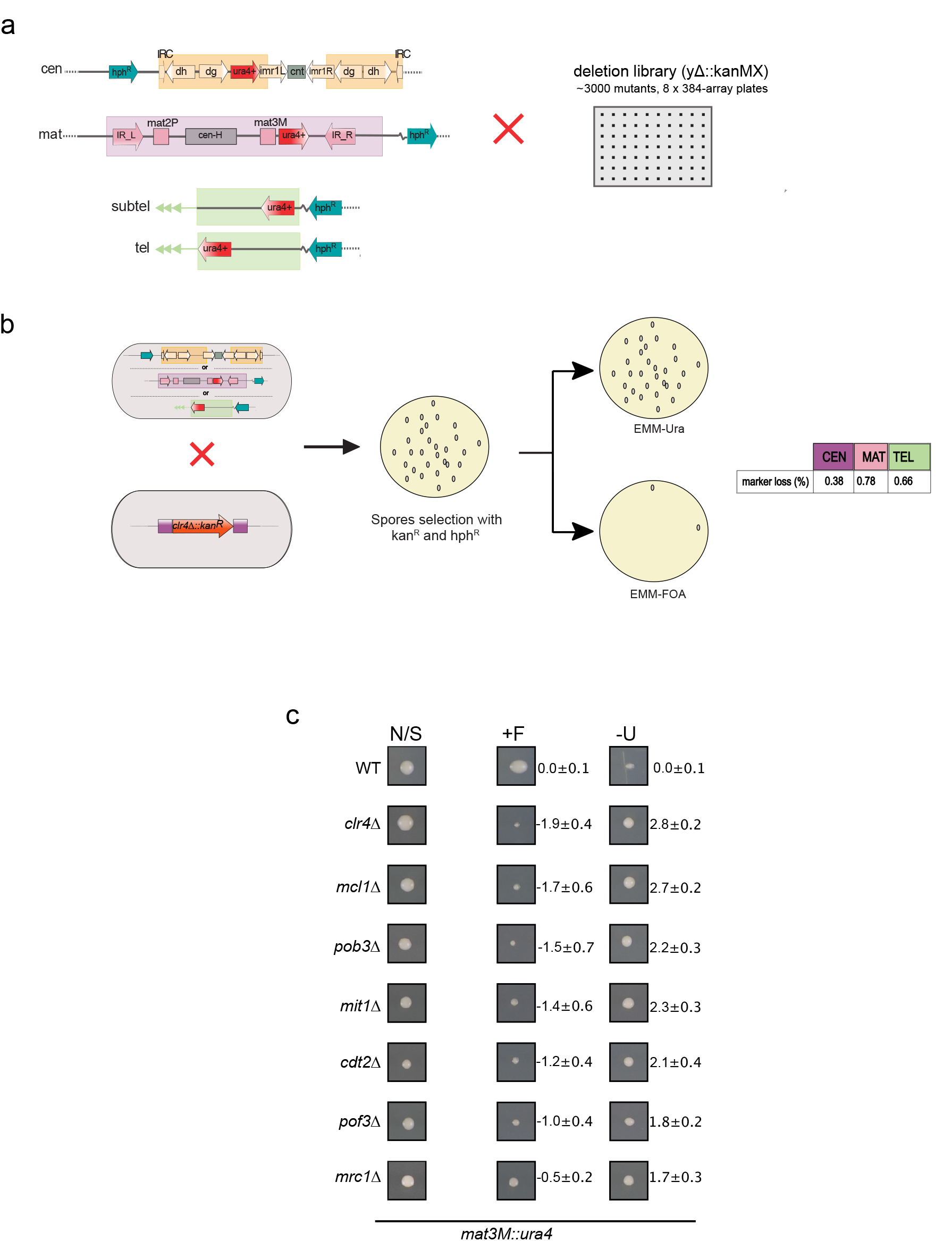
Genetic linkage of selection marker and reproducibility of silencing reporter assays. **a)** Schematic illustration of the *ura4*^*+*^ reporter gene insertions at the four constitutive heterochromatic loci. Shaded areas represent heterochromatin areas, each differentiated by unique color codes. The selection marker (*hphMX6*), which confers hygromycin resistance, was placed 2-4 kb from the heterochromatin boundary within the euchromatin region, ensuring genetic linkage with the *ura4*^*+*^ reporter gene. **b)** Genetic linkage analysis: Yeast strains carrying the *ura4*^*+*^ reporter at various heterochromatic sites (*CEN, MAT, TEL*) were crossed to a *clr4Δ* strain lacking the H3K9 methyltransferase Clr4. Resulting spores that carry both *clr4Δ* and *hphR* marker were selected by replica-plating onto media containing 5-FOA (+F) or media lacking uracil (-U) minimal media to evaluate *ura4*^*+*^ presence. Growth in the presence of 5-FOA indicates a loss of the genetic linkage between *ura4*^*+*^ and *hphR*. Quantification of the genetic linkage tests is shown in the table. **c)** Reproducibility of silencing reporter assays. Displayed are representative colony images from a *MAT* locus screen. The media types are labeled as N/S (non-selective), +F (5-FOA-containing), and -U (uracil-deficient). Quantification of N/S-normalized colony sizes relative to wild-type are shown next to images as log_2_-transformed mean values with standard deviation (SD; (n = 7 biological replicates, each based on two technical replicates).

**Figure S2.**
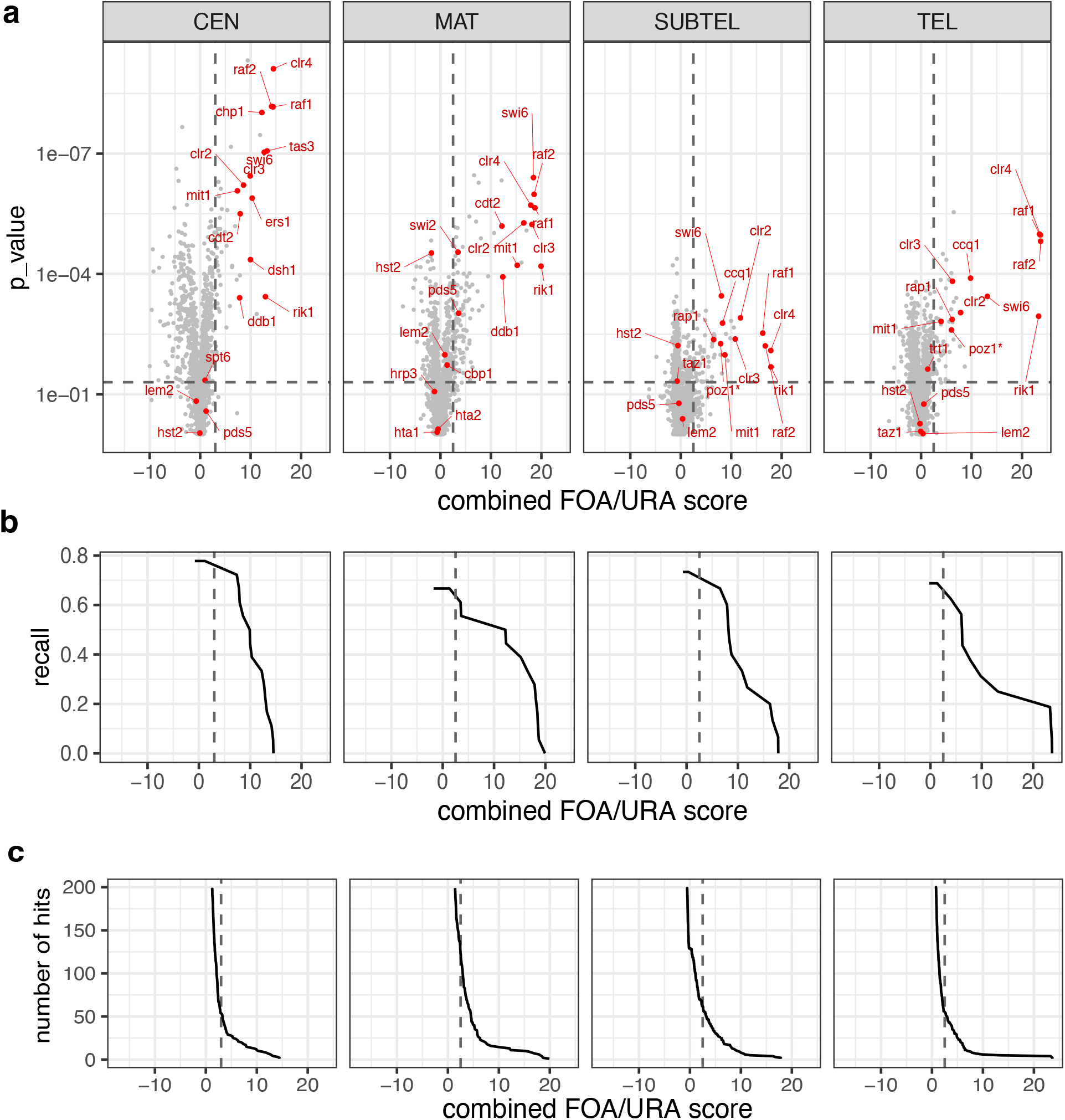
Retrieval of known heterochromatin factors dependent on threshold parameters (recall). **a)** Volcano plots displaying combined FOA/URA scores and reproducibility (P-value from one-sample Student’s t-test) for all mutants from various screens. Prior FOA/URA score combination, relative growth scores for FOA and URA were z-normalized (i.e., setting standard deviation to 1) and median-centered for each heterochromatin reporter, ensuring comparability across all heterochromatic loci (see Methods). Red dots highlight known heterochromatin factors (refer to Table S4). The datasets comprise 3-8 independent biological replicates (*CEN*: 8; *MAT*: 7; *SUBTEL*: 3; *TEL*: 6; each with 2-4 technical replicates). **b)** Recall values of known heterochromatin factors. Plots showing screen sensitivity (ratio of the factors retrieved to the total number of factors) dependent on threshold setting for combined FOA/URA score. **c)** Hit counts relative to threshold settings. The plots show the number of hits dependent on the threshold settings. The dashed lines in (b) and (c) indicate the specific thresholds for the combined FOA/URA scores applied for hit identification: *CEN* = 3; *MAT, SUBTEL, TEL =* 2.5.

**Figure S3.**
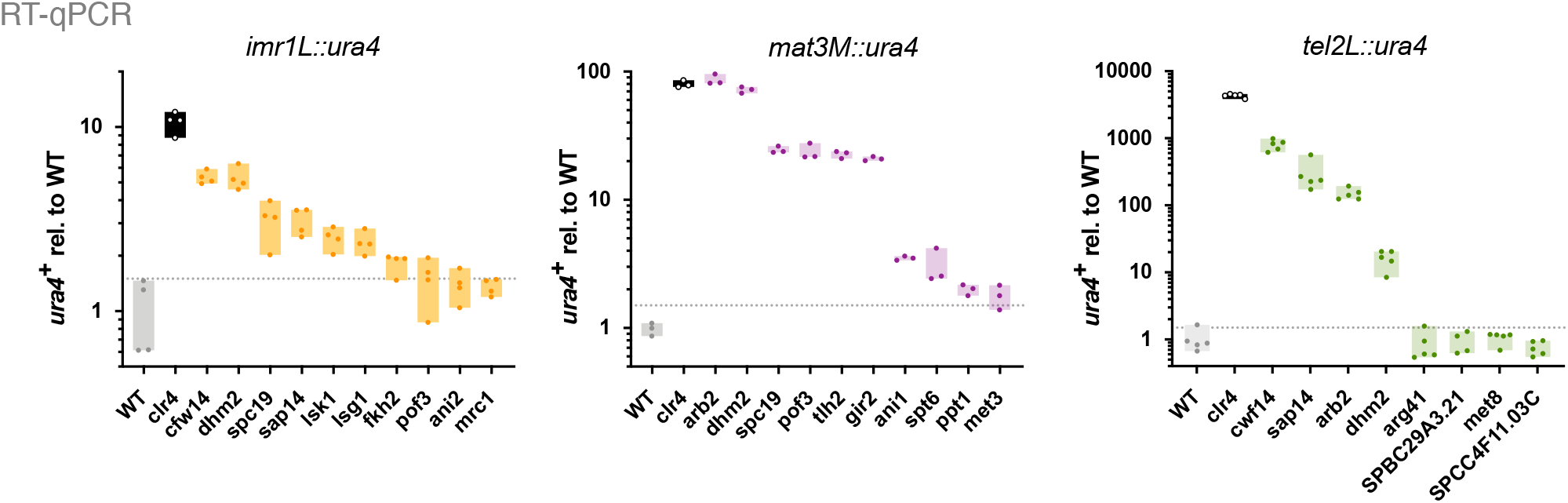
Screen validation by measuring *ura4+* reporter transcript levels (precision). Transcript levels of the *ura4*^*+*^ reporter gene, located at *imr1L, mat3M*, and *tel2L*, were quantified using RT-qPCR. Transcript levels were normalized against *act1* and are depicted relative to the WT mean values (*n* = 4 independent biological replicates). Presented is a representative subset of candidates identified based on the threshold settings employed in this study. The dashed line represents the threshold set for elevated expression (1.5-fold relative to WT). The results are summarized in Suppl. Table S6. Note that for a few candidates identified, *ura4*^+^ transcripts were not found to be increased by RT-qPCR, resulting in precision scores < 1. These deviations might be attributed to differences in the experimental setup (solid minimal media vs. liquid rich media; clonal vs. population-based growth) or in the sensitivity of the assay.

**Figure S4.**
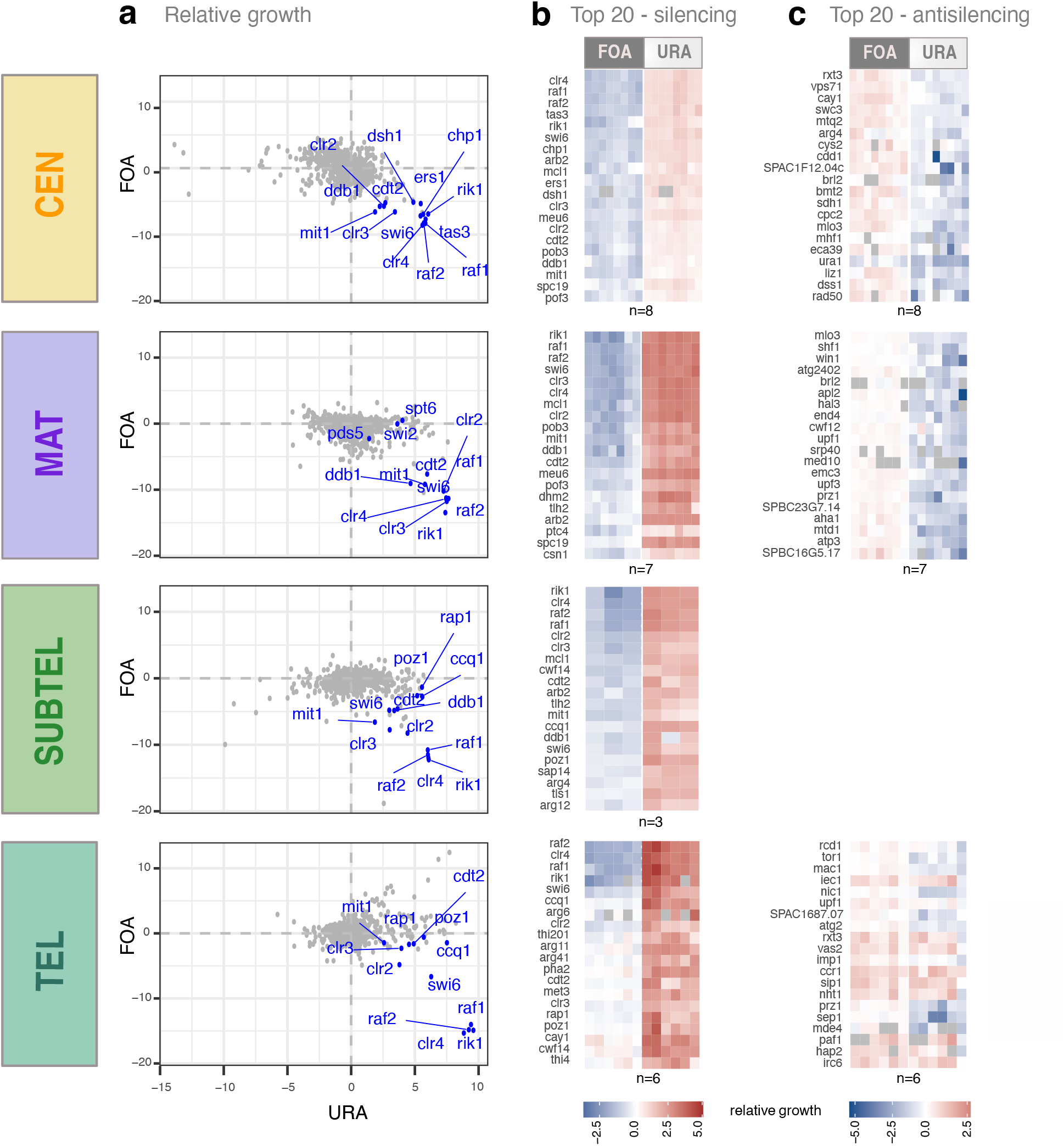
Top candidates among identified silencing and anti-silencing factors. **a)** Scatterplots display log_2_-transformed relative growth values (URA, FOA) for each reporter screen. Median values of 3-8 independent biological replicates (*CEN*: 8; *MAT*: 7; *SUBTEL*: 3; *TEL*: 6; each with 2-4 technical replicates) are shown. Blue dots represent known silencing factors (refer to Suppl. Table S4) detected in the screens. **b-c)** Heatmaps show the relative growth values (log_2_) of the top 20 silencing (b) and anti-silencing (c) candidates. Values are derived from independent biological replicates, with each biological replicate calculated as the average of 2-4 technical replicates.

**Figure S5.**
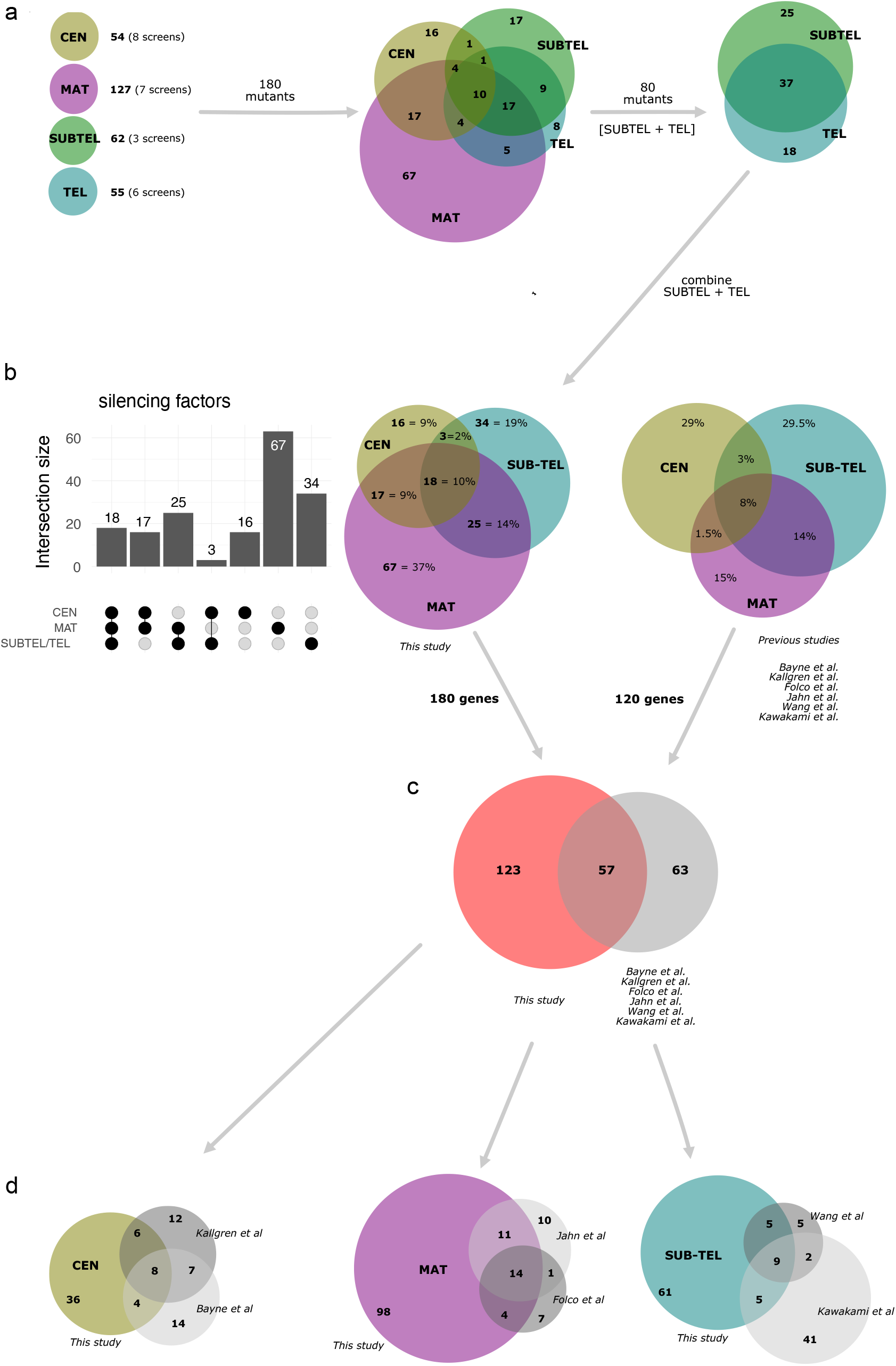
Comparative analysis of silencing hits from this study and previous genome-wide screens. **a)** Venn diagram illustrating the number of silencing hit candidates identified by this study, highlighting the shared defects detected across the four heterochromatic reporter screens. Notably, while overlap between individual reporter screens is limited, a significant overlap is observed between *SUBTEL* and *TEL* screens. **b)** Overlap of hits identified in this study, as shown by Upset plot (right) and Venn diagram (middle), in comparison to previous studies (right). Note, for clearer visualization, candidates from different studies identified by identical or similar (e.g., *SUBTEL, TEL*) reporter systems have been aggregated. **c)** Venn diagram showing the overlap of the total number of hits between this study and previous genome-wide screens. **d)** A series of Venn diagrams illustrate the overlap in domain-specific silencing hits identified in this study compared to those found in previous genome-wide screens.

**Figure S6.**
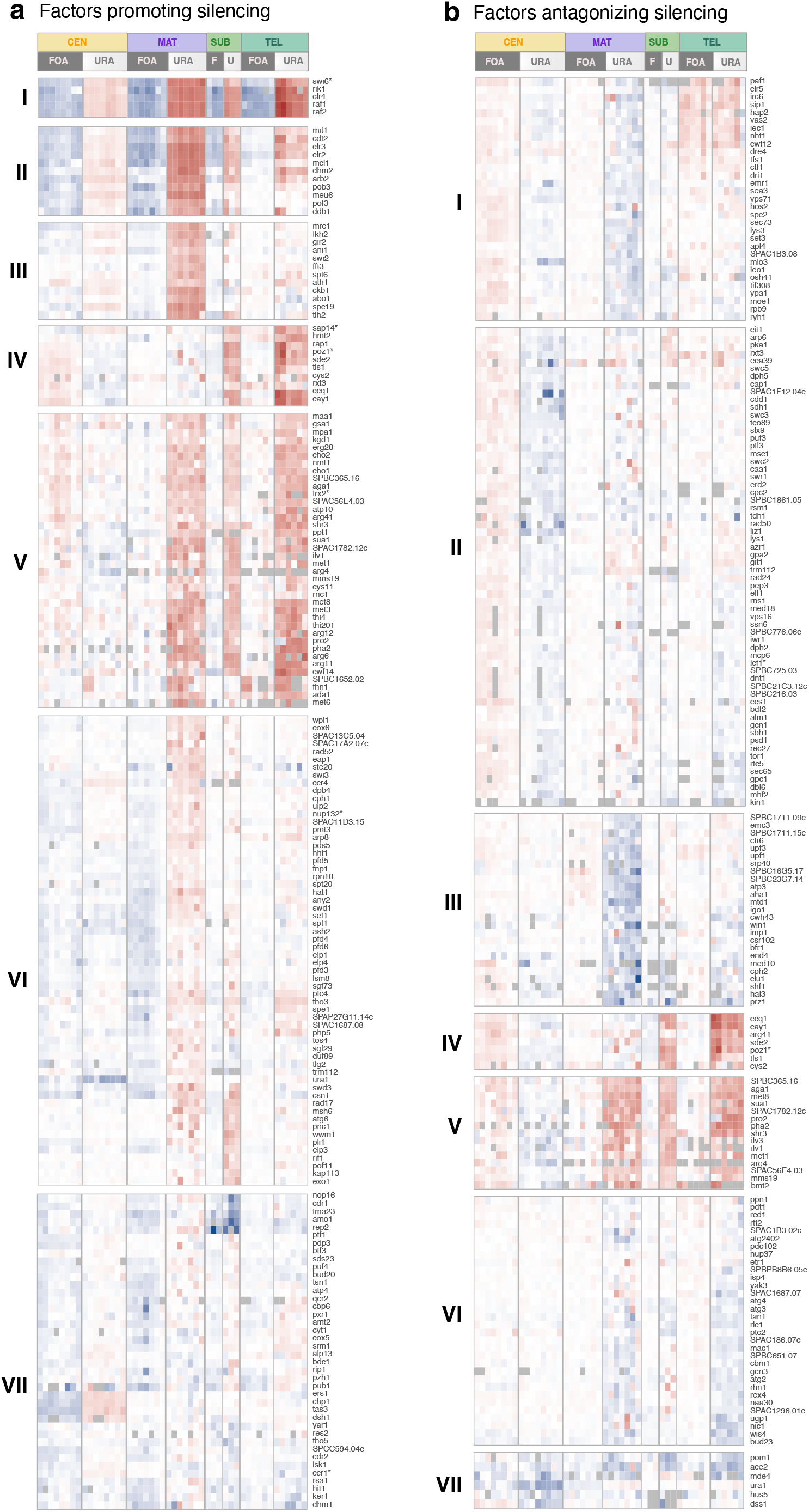
Cluster groups established by k-means clustering of factors promoting or antagonizing silencing. Heatmaps display relative growth values of mutants exhibiting **(a)** significantly reduced and **(b)** significantly enhanced silencing. In this analysis, 176 silencing and 179 anti-silencing mutants were examined. The data represent values derived from 3-8 independent biological replicates (*CEN*: 8; *MAT*: 7; *SUBTEL*: 3; *TEL*: 6; each with 2-4 technical replicates). The gene order within each cluster was determined through subsequent hierarchical clustering (see Methods). Note that 4 silencing mutants (out of 180) and 10 anti-silencing mutants (out of 189) were excluded from the k-means clustering due to missing values. In cases where mutants were mis-annotated in the gene deletion collection, the correct gene name is indicated by an asterisk (see also Suppl. Table S8 and Methods).

**Figure S7.**
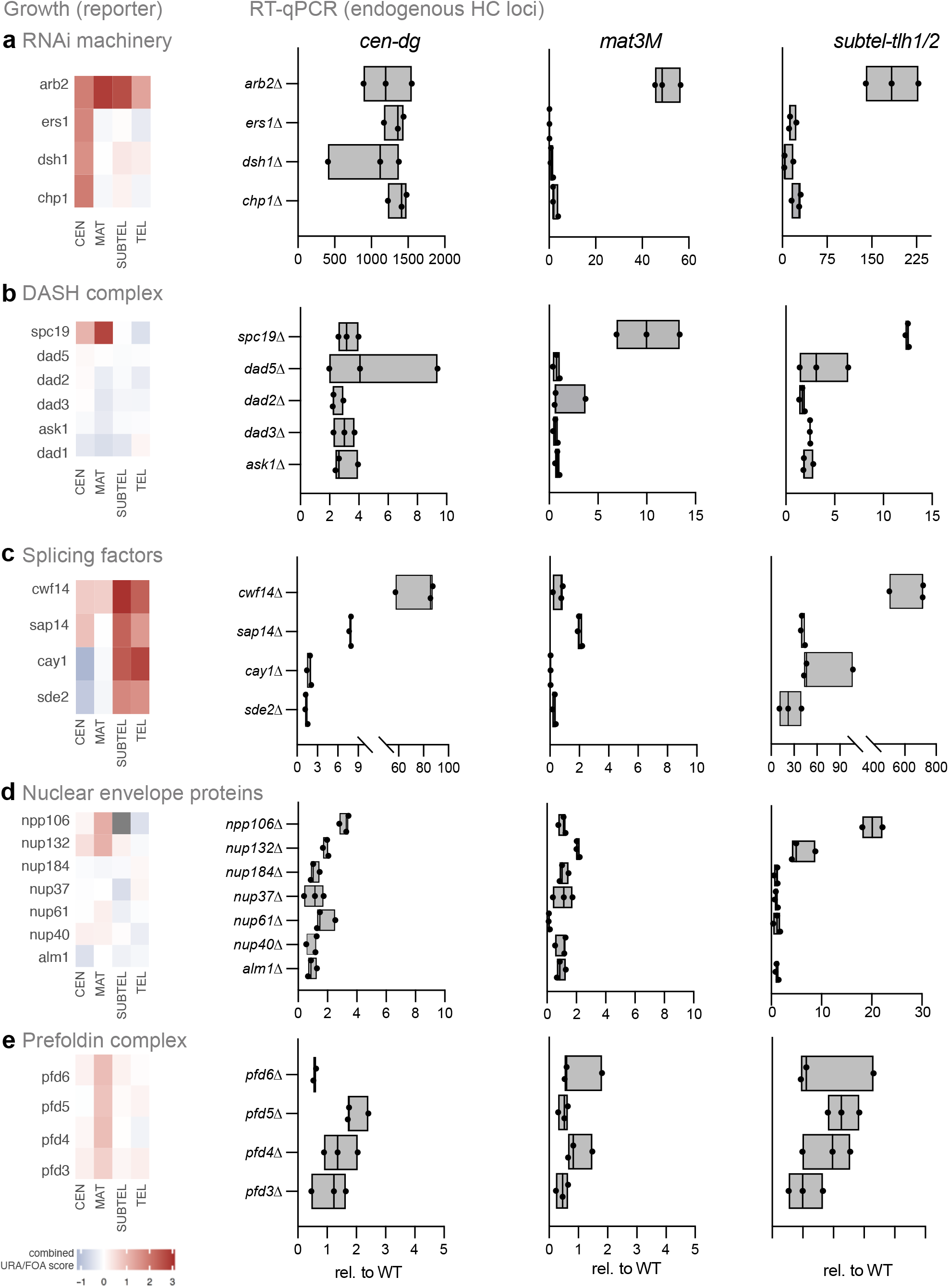
RT-qPCR analysis of mutants associated with specific protein complexes or functional pathways. Reporter growth-based heatmaps and expression analysis of endogenous heterochromatic transcripts in mutants associated with **(a)** RITS, **(b)** DASH, **(c)** splicing factors, **(d)** nuclear envelope protein, and **(e)** prefoldin complex. Heatmaps display median values of combined FOA/URA scores (log_2_). The scores were calculated from 3-8 independent biological replicates (*CEN*: 8; *MAT*: 7; *SUBTEL*: 3; *TEL*: 6; each with 2-4 technical replicates). Plots accompanying each heatmap illustrate the endogenous transcript levels at three heterochromatic loci (*cen-dg, mat3M*, and *tlh1*), as determined by RT-qPCR. Transcript levels, normalized against *act1*, are presented relative to the WT mean values (*n* = 3 independent biological replicates).

**Figure S8.**
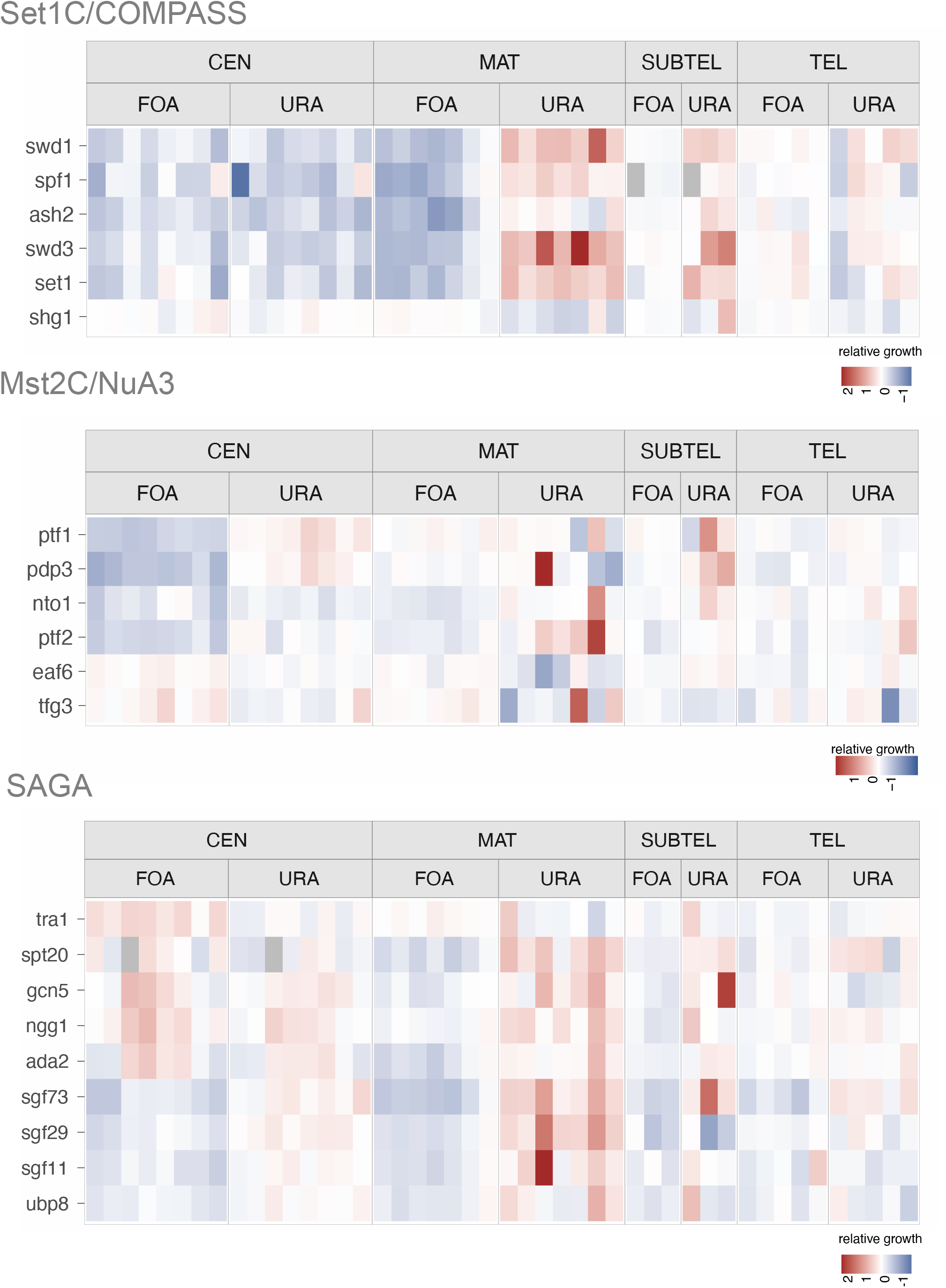
Silencing defects of mutants associated with known chromatin-organizing complexes. Heatmaps show reporter-based relative growth values (log_2_) derived from the genome-wide silencing assays. Mutants associated with a) Set1C/COMPASS, b) Mst2C/NuA3, and c) SAGA complexes are shown.

**Figure S9.**
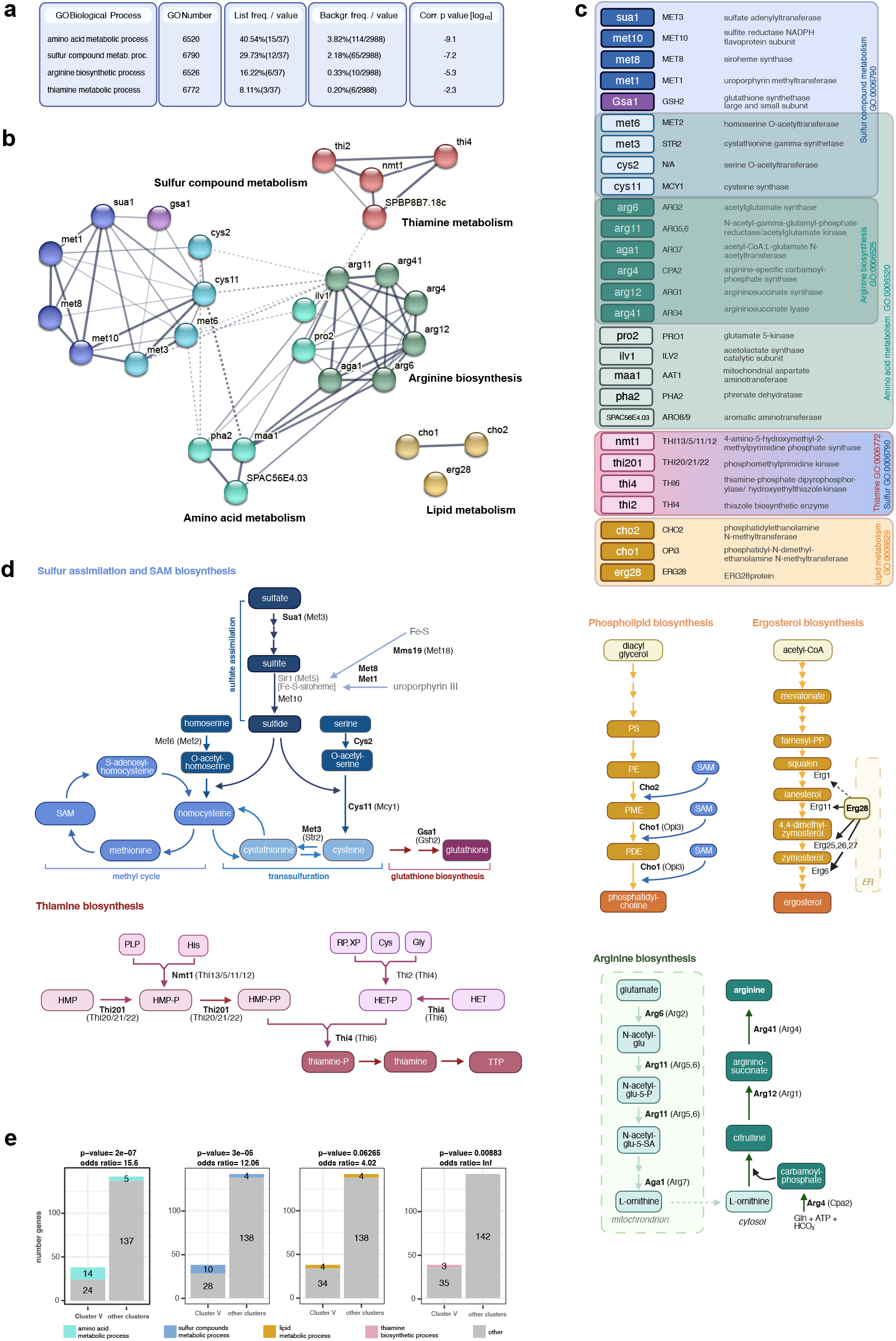
Gene ontology and interaction analysis of silencing factors involved in metabolic processes. **a)** Table presents selected results from gene list enrichment analysi**s** of the 38 genes from silencing factor cluster V. This analysis used the AnGeLi web-based-tool (Bitton et al., 2015) and employed a two-tailed Fisher’s exact test and a false-discovery rate of 0.05 was used for the analysis. **b**) STRING analysis (https://string-db.org/; (Szklarczyk et al., 2023) visualizes interactions among silencing factors from metabolic pathways detailed in (a). **c)** Table list *S. pombe* gene names, along with their *S. cerevisiae* homologs (when available), and provides details on molecular function, metabolic pathways, and associated GO terms. **d**) Schematics illustrating key metabolic pathways in sulfur assimilation/ SAM biosynthesis pathway, phospholipid biosynthesis, ergosterol biosynthesis, thiamine biosynthesis, and arginine biosynthesis. Proteins with bold names represent mutants showing significantly reduced silencing; protein names in brackets indicate *S. cerevisiae* homologs). **e)** Enrichment analysis of the genes involved in metabolic processes across different silencing factor clusters. Bar plots compare Cluster V to other clusters regarding the number of genes associated with distinct metabolic processes, with p-value and odds ratio indicating the significance and strength of association, as determined by Fisher’s exact test.

**Figure S10.**
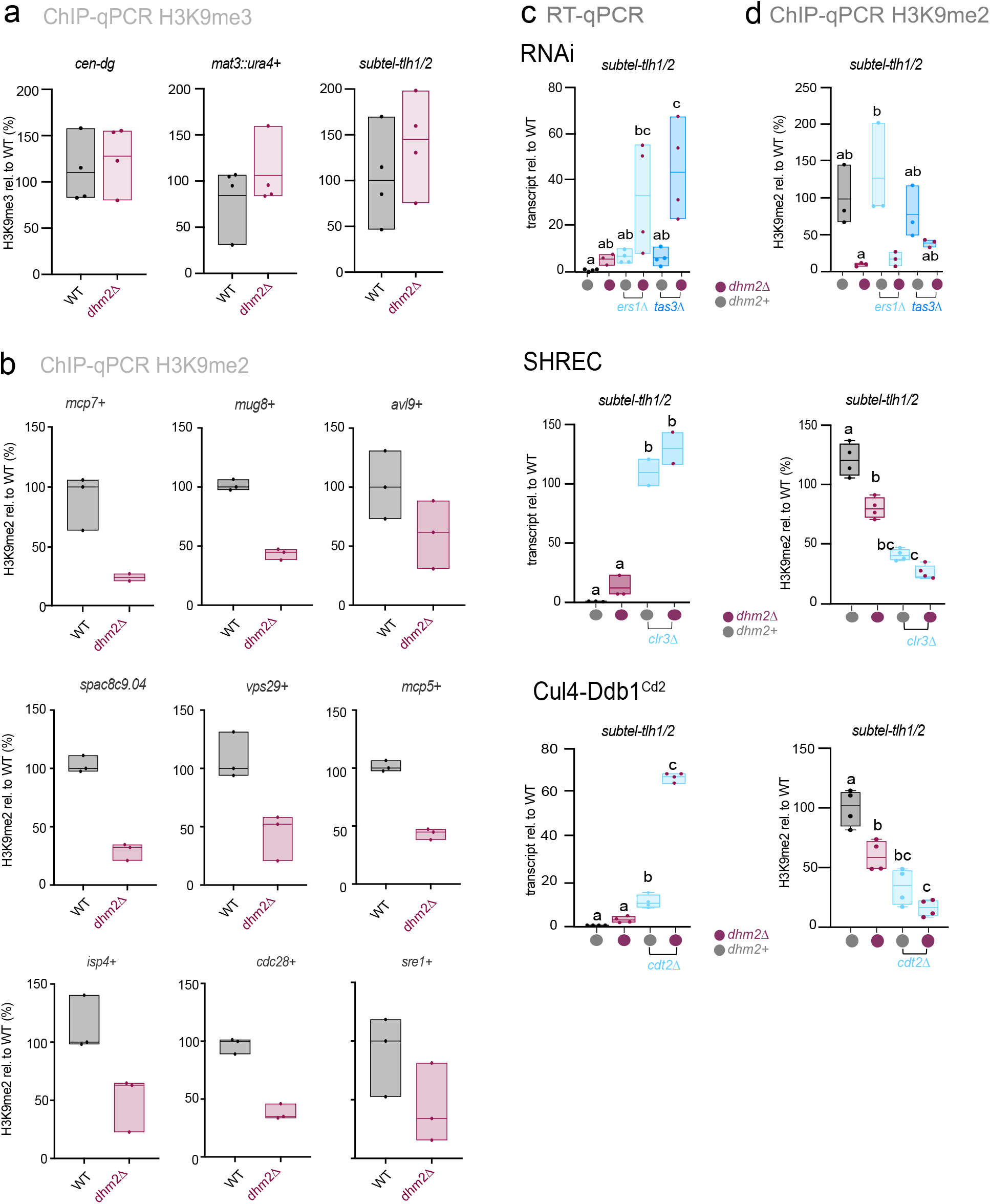
Heterochromatin structure and silencing at constitutive and facultative heterochromatin in *dhm2Δ*. **a**) ChIP-qPCR analysis of H3K9me3 enrichment at *cen-dg* repeats and the *mat3::ura4* reporter gene (*n* = 3-4 independent biological replicates). **b**) ChIP-qPCR analysis of H3K9me2 levels of facultative heterochromatin islands (*n* = 3 independent biological replicates). **c**) RT-qPCR quantification of *tlh1* transcript levels quantified by RT-qPCR in the indicated strains (*n* = 3-4 independent biological replicates). Data are normalized to *act1* transcript level and presented relative to WT median value. **d)** ChIP-qPCR analysis of H3K9me2 levels at *tlh1*^*+*^ in the indicated strains (*n* = 3-4). For ChIP analysis in (a), (b), and (c), immunoprecipitated (IP) samples, normalized to input, are further standardized against the average of two euchromatic loci (*act1*^*+*^ and *tef3*^*+*^). Data are presented relative to the WT median value. Statistical analysis for (c) and (d) employed one-way ANOVA, with Tukey’s *post hoc* test identifying significant differences at *P* < 0.05. Letters denote groups with significant differences.

**Figure S11.**
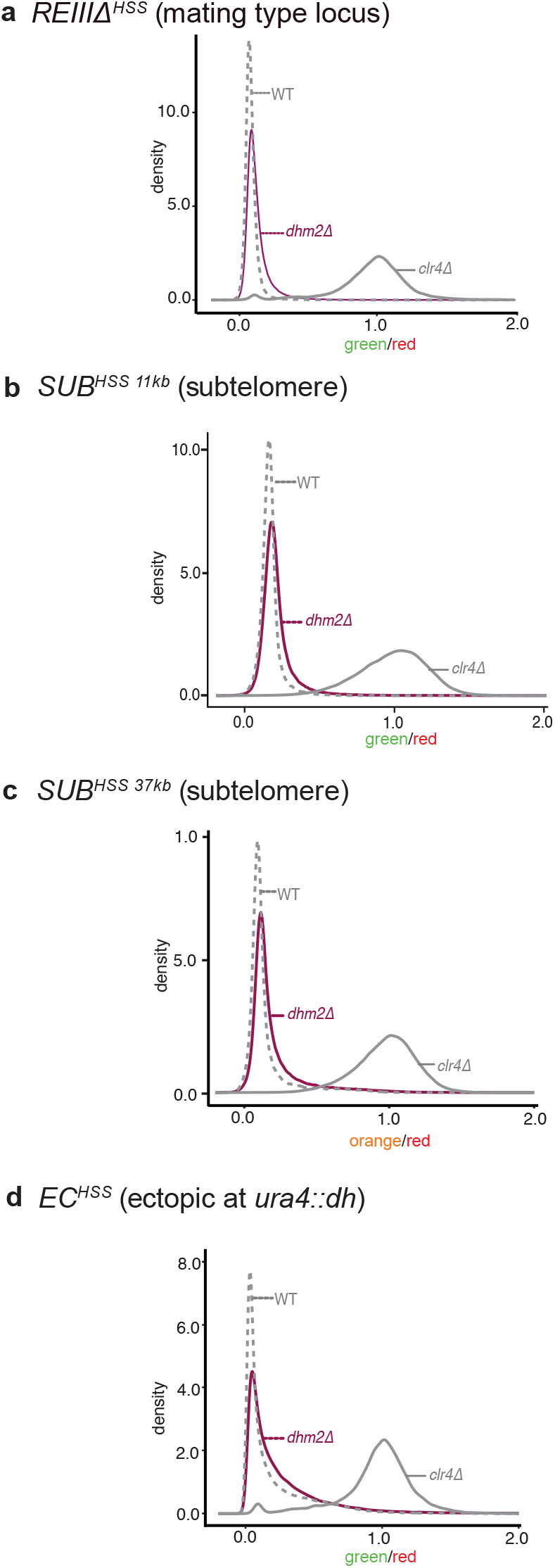
Assessment of heterochromatin establishment at single-cell level at various nucleation sites in *dhm2Δ*. Density plots of flow cytometry measurements illustrating reporter expression in wild-type (WT; dashed line), *dhm2Δ* (maroon solid line), and *clr4Δ* (gray solid line) strains across various heterochromatin domains: **a)** *cenH* region of the mating type locus (ΔREIII^HSS^). **b)** subtelomeric locus 11 kb downstream of the telomeric repeats (*SUB*^*HSS 11kb*^). **c)** Subtelomeric locus 37 kb downstream of the telomeric repeats (*SUB*^*HSS 37kb*^). **d)** Ectopic locus created by inserting the pericentromeric *dh* element next to endogenous *ura4*^*+*^ locus (*EC*^*HSS*^). The x-axis displays relative expression values of the SF-GFPsp (green) or the mKO2sp (orange) reporter. The y-axis represents the density of the cell population. Reporter expression is normalized to E2C (red; noise filter) expressed from an adjacent locus located in euchromatin and is shown relative to the median expression in the *clr4Δ* strain.

**Figure S12.**
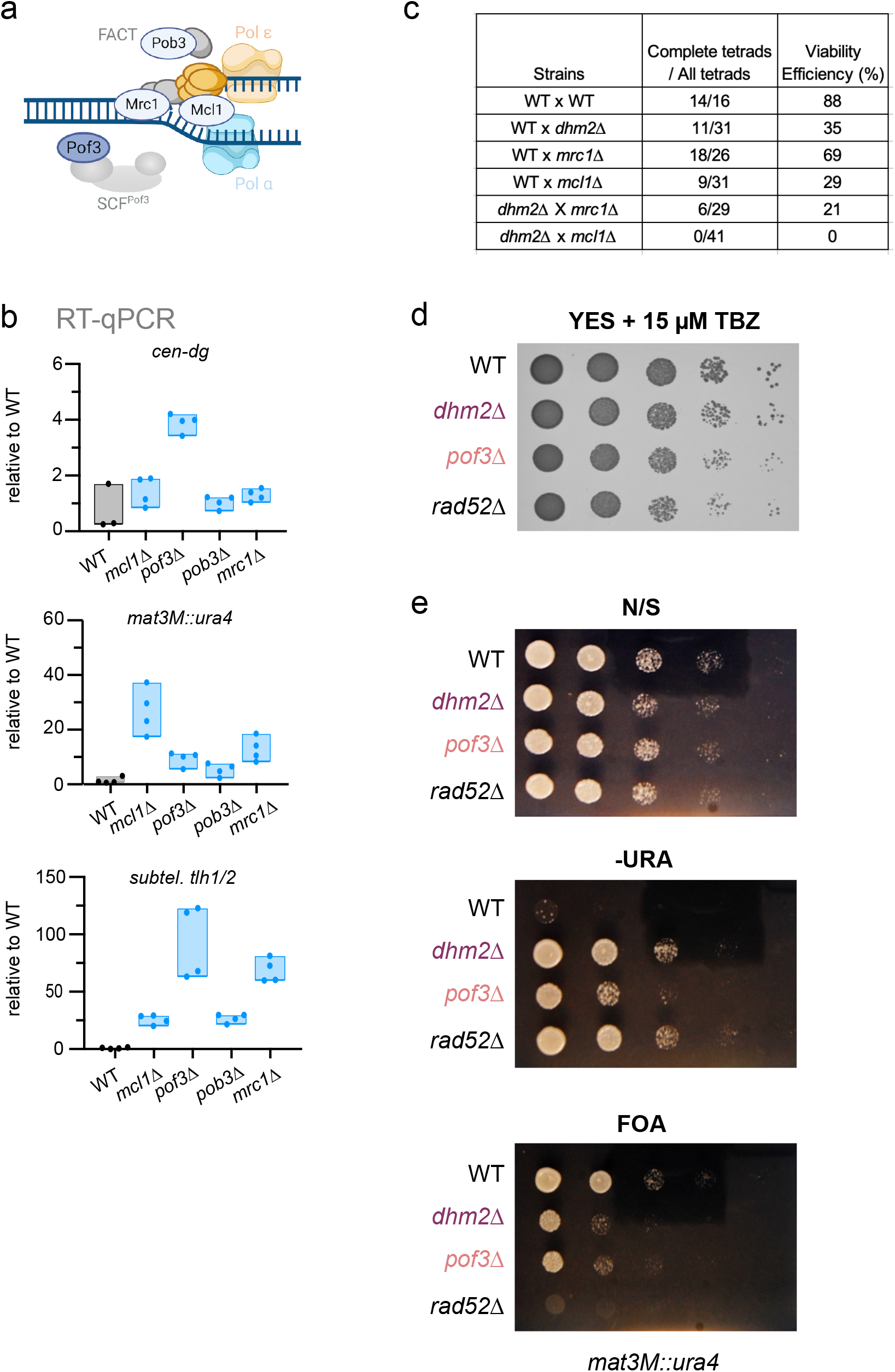
Heterochromatin silencing in various mutants associated with DNA replication. **a)** Schematic illustrating factors involved in DNA replication. **b)** RT-qPCR quantification of heterochromatic transcripts (*cen-dg, mat3::ura4* and *tlh1/2*) in mutants lacking Mcl1^Ctf4^, Pof3^F-box^, Pob3^FACT^ and Mrc1^Claspin^. Data are normalized to *act1* transcript level and presented relative to the WT median value (n = 4). **c)** Viability efficiency (%) was determined by counting the number of complete tetrad (four-spored ascus) from the representative strains **d)** Sensitivity toward thiabendazole (TBZ). Tenfold serial dilutions of the indicated strains were plated on YES medium supplemented with 15 μM TBZ and incubated for 3 days at 32°C. **e**) Reporter silencing assay. Tenfold serial dilutions of indicated strains were plated on EMM (minimal growth medium) under various conditions (N/S, non-selective; -URA, without uracil; FOA, supplemented with fluoroorotic acid) and incubated for 3 days at 32°C.

## Notes

https://github.com/zsarkadi/Muhammad-Sarkadi-et-al.

### Competing Interest Statement

The authors have declared no competing interest.

## References

Allshire, R.C., Ekwall, K., 2015. Epigenetic Regulation of Chromatin States in Schizosaccharomyces pombe. Cold Spring Harb. Perspect. Biol. 7. 10.1101/cshperspect.a018770

Allshire, R.C., Madhani, H.D., 2018. Ten principles of heterochromatin formation and function. Nat. Rev. Mol. Cell Biol. 19, 229–244. 10.1038/nrm.2017.119

Allshire, R.C., Nimmo, E.R., Ekwall, K., Javerzat, J.P., Cranston, G., 1995. Mutations derepressing silent centromeric domains in fission yeast disrupt chromosome segregation. Genes Dev. 9, 218–233. 10.1101/gad.9.2.218

Al-Sady, B., Greenstein, R.A., El-Samad, H.J., Braun, S., Madhani, H.D., 2016. Sensitive and Quantitative Three-Color Protein Imaging in Fission Yeast Using Spectrally Diverse, Recoded Fluorescent Proteins with Experimentally-Characterized In Vivo Maturation Kinetics. PLOS ONE 11, e0159292. 10.1371/journal.pone.0159292

Anil, A.T., Choudhary, K., Pandian, R., Gupta, P., Thakran, P., Singh, A., Sharma, M., Mishra, S.K., 2022. Splicing of branchpoint-distant exons is promoted by Cactin, Tls1 and the ubiquitin-fold-activated Sde2. Nucleic Acids Res. 50, 10000–10014. 10.1093/nar/gkac769

Audergon, P.N.C.B., Catania, S., Kagansky, A., Tong, P., Shukla, M., Pidoux, A.L., Allshire, R.C., 2015. Epigenetics. Restricted epigenetic inheritance of H3K9 methylation. Science 348, 132–135. 10.1126/science.1260638

Ayoub, N., Noma, K., Isaac, S., Kahan, T., Grewal, S.I.S., Cohen, A., 2003. A novel jmjC domain protein modulates heterochromatization in fission yeast. Mol. Cell. Biol. 23, 4356–4370. 10.1128/MCB.23.12.4356-4370.2003

Bannister, A.J., Zegerman, P., Partridge, J.F., Miska, E.A., Thomas, J.O., Allshire, R.C., Kouzarides, T., 2001. Selective recognition of methylated lysine 9 on histone H3 by the HP1 chromo domain. Nature 410, 120–124. 10.1038/35065138

Barrales, R.R., Forn, M., Georgescu, P.R., Sarkadi, Z., Braun, S., 2016. Control of heterochromatin localization and silencing by the nuclear membrane protein Lem2. Genes Dev. 30, 133–148. 10.1101/gad.271288.115

Baryshnikova, A., Costanzo, M., Dixon, S., Vizeacoumar, F.J., Myers, C.L., Andrews, B., Boone, C., 2010. Chapter 7 - Synthetic Genetic Array (SGA) Analysis in Saccharomyces cerevisiae and Schizosaccharomyces pombe, in: Methods in Enzymology, Guide to Yeast Genetics: Functional Genomics, Proteomics, and Other Systems Analysis. Academic Press, pp. 145–179. 10.1016/S0076-6879(10)70007-0

Bayne, E.H., Bijos, D.A., White, S.A., de Lima Alves, F., Rappsilber, J., Allshire, R.C., 2014. A systematic genetic screen identifies new factors influencing centromeric heterochromatin integrity in fission yeast. Genome Biol. 15, 481. 10.1186/s13059-014-0481-4

Bayne, E.H., White, S.A., Kagansky, A., Bijos, D.A., Sanchez-Pulido, L., Hoe, K.-L., Kim, D.-U., Park, H.-O., Ponting, C.P., Rappsilber, J., Allshire, R.C., 2010. Stc1: a critical link between RNAi and chromatin modification required for heterochromatin integrity. Cell 140, 666–677. 10.1016/j.cell.2010.01.038

Bitton, D.A., Schubert, F., Dey, S., Okoniewski, M., Smith, G.C., Khadayate, S., Pancaldi, V., Wood, V., Bähler, J., 2015. AnGeLi: A Tool for the Analysis of Gene Lists from Fission Yeast. Front. Genet. 6, 330. 10.3389/fgene.2015.00330

Braun, S., Garcia, J.F., Rowley, M., Rougemaille, M., Shankar, S., Madhani, H.D., 2011. The Cul4-Ddb1(Cdt)2 ubiquitin ligase inhibits invasion of a boundary-associated antisilencing factor into heterochromatin. Cell 144, 41–54. 10.1016/j.cell.2010.11.051

Cooper, J.P., Nimmo, E.R., Allshire, R.C., Cech, T.R., 1997. Regulation of telomere length and function by a Myb-domain protein in fission yeast. Nature 385, 744–747. 10.1038/385744a0

Cutter DiPiazza, A.R., Taneja, N., Dhakshnamoorthy, J., Wheeler, D., Holla, S., Grewal, S.I.S., 2021. Spreading and epigenetic inheritance of heterochromatin require a critical density of histone H3 lysine 9 tri-methylation. Proc. Natl. Acad. Sci. U. S. A. 118, e2100699118. 10.1073/pnas.2100699118

Deng, X., Zhou, H., Zhang, G., Wang, W., Mao, L., Zhou, X., Yu, Y., Lu, H., 2015. Sgf73, a subunit of SAGA complex, is required for the assembly of RITS complex in fission yeast. Sci. Rep. 5, 14707. 10.1038/srep14707

Egan, E.D., Braun, C.R., Gygi, S.P., Moazed, D., 2014. Post-transcriptional regulation of meiotic genes by a nuclear RNA silencing complex. RNA N. Y. N 20, 867–881. 10.1261/rna.044479.114

Ekwall, K., Cranston, G., Allshire, R.C., 1999. Fission yeast mutants that alleviate transcriptional silencing in centromeric flanking repeats and disrupt chromosome segregation. Genetics 153, 1153–1169. 10.1093/genetics/153.3.1153

Fan, J., Krautkramer, K.A., Feldman, J.L., Denu, J.M., 2015. Metabolic Regulation of Histone Post-Translational Modifications. ACS Chem. Biol. 10, 95–108. 10.1021/cb500846u

Flury, V., Georgescu, P.R., Iesmantavicius, V., Shimada, Y., Kuzdere, T., Braun, S., Bühler, M., 2017. The Histone Acetyltransferase Mst2 Protects Active Chromatin from Epigenetic Silencing by Acetylating the Ubiquitin Ligase Brl1. Mol. Cell 67, 294–307.e9. 10.1016/j.molcel.2017.05.026

Folco, H.D., McCue, A., Balachandran, V., Grewal, S.I.S., 2019. Cohesin Impedes Heterochromatin Assembly in Fission Yeast Cells Lacking Pds5. Genetics 213, 127–141. 10.1534/genetics.119.302256

Gal, C., Murton, H.E., Subramanian, L., Whale, A.J., Moore, K.M., Paszkiewicz, K., Codlin, S., Bähler, J., Creamer, K.M., Partridge, J.F., Allshire, R.C., Kent, N.A., Whitehall, S.K., 2016. Abo1, a conserved bromodomain AAA-ATPase, maintains global nucleosome occupancy and organisation. EMBO Rep. 17, 79–93. 10.15252/embr.201540476

Gari, K., León Ortiz, A.M., Borel, V., Flynn, H., Skehel, J.M., Boulton, S.J., 2012. MMS19 links cytoplasmic iron-sulfur cluster assembly to DNA metabolism. Science 337, 243–245. 10.1126/science.1219664

Georgescu, P.R., Capella, M., Fischer-Burkart, S., Braun, S., 2020. The euchromatic histone mark H3K36me3 preserves heterochromatin through sequestration of an acetyltransferase complex in fission yeast. Microb. Cell 7, 80–92. 10.15698/mic2020.03.711

Gerace, E.L., Halic, M., Moazed, D., 2010. The methyltransferase activity of Clr4Suv39h triggers RNAi independently of histone H3K9 methylation. Mol. Cell 39, 360–372. 10.1016/j.mol-cel.2010.07.017

Greenstein, R.A., Jones, S.K., Spivey, E.C., Rybarski, J.R., Finkelstein, I.J., Al-Sady, B., 2018. Noncoding RNA-nucleated heterochromatin spreading is intrinsically labile and requires accessory elements for epigenetic stability. eLife 7, e32948. 10.7554/eLife.32948

Greenstein, R.A., Ng, H., Barrales, R.R., Tan, C., Braun, S., Al-Sady, B., 2022. Local chromatin context regulates the genetic requirements of the heterochromatin spreading reaction. PLoS Genet. 18, e1010201. 10.1371/journal.pgen.1010201

Grewal, S.I.S., 2023. The molecular basis of heterochromatin assembly and epigenetic inheritance. Mol. Cell 83, 1767–1785. 10.1016/j.molcel.2023.04.020

Han, T.X., Xu, X.-Y., Zhang, M.-J., Peng, X., Du, L.-L., 2010. Global fitness profiling of fission yeast deletion strains by barcode sequencing. Genome Biol. 11, R60. 10.1186/gb-2010-11-6-r60

Hansen, K.R., Burns, G., Mata, J., Volpe, T.A., Martienssen, R.A., Bähler, J., Thon, G., 2005. Global effects on gene expression in fission yeast by silencing and RNA interference machineries. Mol. Cell. Biol. 25, 590–601. 10.1128/MCB.25.2.590-601.2005

Hansen, K.R., Ibarra, P.T., Thon, G., 2006. Evolutionary-conserved te-lomere-linked helicase genes of fission yeast are repressed by silencing factors, RNAi components and the telomere-binding protein Taz1. Nucleic Acids Res. 34, 78–88. 10.1093/nar/gkj415

Harr, J.C., Gonzalez-Sandoval, A., Gasser, S.M., 2016. Histones and histone modifications in perinuclear chromatin anchoring: from yeast to man. EMBO Rep. 17, 139–155. 10.15252/embr.201541809

Hayashi, A., Ishida, M., Kawaguchi, R., Urano, T., Murakami, Y., Nakayama, J., 2012. Heterochromatin protein 1 homologue Swi6 acts in concert with Ers1 to regulate RNAi-directed heterochromatin assembly. Proc. Natl. Acad. Sci. U. S. A. 109, 6159–6164. 10.1073/pnas.1116972109

Hayashi, T., Teruya, T., Chaleckis, R., Morigasaki, S., Yanagida, M., 2018. S-Adenosylmethionine Synthetase Is Required for Cell Growth, Maintenance of G0 Phase, and Termination of Quiescence in Fission Yeast. iScience 5, 38–51. 10.1016/j.isci.2018.06.011

Hirano, Y., Kinugasa, Y., Kubota, Y., Obuse, C., Haraguchi, T., Hiraoka, Y., 2023. Inner nuclear membrane proteins Lem2 and Bqt4 interact with different lipid synthesis enzymes in fission yeast. J. Biochem. (Tokyo) 174, 33–46. 10.1093/jb/mvad017

Holla, S., Dhakshnamoorthy, J., Folco, H.D., Balachandran, V., Xiao, H., Sun, L., Wheeler, D., Zofall, M., Grewal, S.I.S., 2020. Positioning Heterochromatin at the Nuclear Periphery Suppresses Histone Turnover to Promote Epigenetic Inheritance. Cell 180, 150–164.e15. 10.1016/j.cell.2019.12.004

Holoch, D., Moazed, D., 2015. RNA-mediated epigenetic regulation of gene expression. Nat. Rev. Genet. 16, 71–84. 10.1038/nrg3863

Hong, E.-J.E., Villén, J., Gerace, E.L., Gygi, S.P., Moazed, D., 2005. A cullin E3 ubiquitin ligase complex associates with Rik1 and the Clr4 histone H3-K9 methyltransferase and is required for RNAi-mediated heterochromatin formation. RNA Biol. 2, 106–111. 10.4161/rna.2.3.2131

Horn, P.J., Bastie, J.-N., Peterson, C.L., 2005. A Rik1-associated, cullin-dependent E3 ubiquitin ligase is essential for heterochromatin formation. Genes Dev. 19, 1705–1714. 10.1101/gad.1328005

Iglesias, N., Paulo, J.A., Tatarakis, A., Wang, X., Edwards, A.L., Bhanu, N.V., Garcia, B.A., Haas, W., Gygi, S.P., Moazed, D., 2020. Native Chromatin Proteomics Reveals a Role for Specific Nucleoporins in Heterochromatin Organization and Maintenance. Mol. Cell 77, 51–66.e8. 10.1016/j.mol-cel.2019.10.018

Jahn, L.J., Mason, B., Brøgger, P., Toteva, T., Nielsen, D.K., Thon, G., 2018. Dependency of Heterochromatin Domains on Replication Factors. G3 Bethesda Md 8, 477–489. 10.1534/g3.117.300341

Jain, R., Vanamee, E.S., Dzikovski, B.G., Buku, A., Johnson, R.E., Prakash, L., Prakash, S., Aggarwal, A.K., 2014. An iron-sulfur cluster in the polymerase domain of yeast DNA polymerase ε. J. Mol. Biol. 426, 301–308. 10.1016/j.jmb.2013.10.015

Janke, C., Magiera, M.M., Rathfelder, N., Taxis, C., Reber, S., Maekawa, H., Moreno-Borchart, A., Doenges, G., Schwob, E., Schiebel, E., Knop, M., 2004. A versatile toolbox for PCR-based tagging of yeast genes: new fluorescent proteins, more markers and promoter substitution cassettes. Yeast 21, 947–962. 10.1002/yea.1142

Jia, S., Kobayashi, R., Grewal, S.I.S., 2005. Ubiquitin ligase component Cul4 associates with Clr4 histone methyltransferase to assemble heterochromatin. Nat. Cell Biol. 7, 1007–1013. 10.1038/ncb1300

Jia, S., Noma, K., Grewal, S.I.S., 2004. RNAi-independent heterochromatin nucleation by the stress-activated ATF/CREB family proteins. Science 304, 1971–1976. 10.1126/sci-ence.1099035

Job, G., Brugger, C., Xu, T., Lowe, B.R., Pfister, Y., Qu, C., Shanker, S., Baños Sanz, J.I., Partridge, J.F., Schalch, T., 2016. SHREC Silences Heterochromatin via Distinct Remodeling and Deacetylation Modules. Mol. Cell 62, 207–221. 10.1016/j.mol-cel.2016.03.016

Kallgren, S.P., Andrews, S., Tadeo, X., Hou, H., Moresco, J.J., Tu, P.G., Yates, J.R., Nagy, P.L., Jia, S., 2014. The proper splicing of RNAi factors is critical for pericentric heterochromatin assembly in fission yeast. PLoS Genet. 10, e1004334. 10.1371/journal.pgen.1004334

Kanoh, J., Sadaie, M., Urano, T., Ishikawa, F., 2005. Telomere binding protein Taz1 establishes Swi6 heterochromatin independently of RNAi at telomeres. Curr. Biol. CB 15, 1808–1819. 10.1016/j.cub.2005.09.041

Kassube, S.A., Thomä, N.H., 2020. Structural insights into Fe-S protein biogenesis by the CIA targeting complex. Nat. Struct. Mol. Biol. 27, 735–742. 10.1038/s41594-020-0454-0

Kawakami, K., Hayashi, A., Nakayama, J.-I., Murakami, Y., 2012. A novel RNAi protein, Dsh1, assembles RNAi machinery on chromatin to amplify heterochromatic siRNA. Genes Dev. 26, 1811–1824. 10.1101/gad.190272.112

Kawakami, K., Ueno, Y., Hayama, N., Tanaka, K., 2023. Mrc1Claspin is essential for heterochromatin maintenance in Schizosaccharomyces pombe. 10.1101/2023.03.28.534615

Kiely, C.M., Marguerat, S., Garcia, J.F., Madhani, H.D., Bähler, J., Winston, F., 2011. Spt6 Is Required for Heterochromatic Silencing in the Fission Yeast Schizosaccharomyces pombe. Mol. Cell. Biol. 31, 4193–4204. 10.1128/MCB.05568-11

Kim, H.S., Choi, E.S., Shin, J.A., Jang, Y.K., Park, S.D., 2004. Regulation of Swi6/HP1-dependent heterochromatin assembly by cooperation of components of the mitogen-activated protein kinase pathway and a histone deacetylase Clr6. J. Biol. Chem. 279, 42850– 42859. 10.1074/jbc.M407259200

Kinugasa, Y., Hirano, Y., Sawai, M., Ohno, Y., Shindo, T., Asakawa, H., Chikashige, Y., Shibata, S., Kihara, A., Haraguchi, T., Hiraoka, Y., 2019. The very-long-chain fatty acid elongase Elo2 rescues lethal defects associated with loss of the nuclear barrier function in fission yeast cells. J. Cell Sci. 132, jcs229021. 10.1242/jcs.229021

Kowalik, K.M., Shimada, Y., Flury, V., Stadler, M.B., Batki, J., Bühler, M., 2015. The Paf1 complex represses small-RNA-mediated epigenetic gene silencing. Nature 520, 248–252. 10.1038/nature14337

Larsson, J., Zhang, J., Rasmuson-Lestander, A., 1996. Mutations in the Drosophila melanogaster gene encoding S-adenosylmethionine synthetase [corrected] suppress position-effect variegation. Genetics 143, 887–896. 10.1093/genetics/143.2.887

Lee, N.N., Chalamcharla, V.R., Reyes-Turcu, F., Mehta, S., Zofall, M., Balachandran, V., Dhakshnamoorthy, J., Taneja, N., Yamanaka, S., Zhou, M., Grewal, S.I.S., 2013. Mtr4-like protein coordinates nuclear RNA processing for heterochromatin assembly and for telomere maintenance. Cell 155, 1061–1074. 10.1016/j.cell.2013.10.027

Li, F., Goto, D.B., Zaratiegui, M., Tang, X., Martienssen, R., Cande, W.Z., 2005. Two novel proteins, dos1 and dos2, interact with rik1 to regulate heterochromatic RNA interference and histone modification. Curr. Biol. CB 15, 1448–1457. 10.1016/j.cub.2005.07.021

Li, F., Martienssen, R., Cande, W.Z., 2011. Coordination of DNA Replication and Histone Modification by the Rik1-Dos2 Complex. Nature 475, 244–248. 10.1038/nature10161

Lim, K.K., Teo, H.Y., Tan, Y.Y., Zeng, Y.B., Lam, U.T.F., Choolani, M., Chen, E.S., 2021. Fission Yeast Methylenetetrahydrofolate Reductase Ensures Mitotic and Meiotic Chromosome Segregation Fidelity. Int. J. Mol. Sci. 22, 639. 10.3390/ijms22020639

Lisby, M., Rothstein, R., Mortensen, U.H., 2001. Rad52 forms DNA repair and recombination centers during S phase. Proc. Natl. Acad. Sci. U. S. A. 98, 8276–8282. 10.1073/pnas.121006298

Maculins, T., Nkosi, P.J., Nishikawa, H., Labib, K., 2015. Tethering of SCF(Dia2) to the Replisome Promotes Efficient Ubiquitylation and Disassembly of the CMG Helicase. Curr. Biol. CB 25, 2254–2259. 10.1016/j.cub.2015.07.012

Mamnun, Y.M., Katayama, S., Toda, T., 2006. Fission yeast Mcl1 interacts with SCFPof3 and is required for centromere formation. Bi-ochem. Biophys. Res. Commun. 350, 125–130. 10.1016/j.bbrc.2006.09.024

Martín Caballero, L., Capella, M., Barrales, R.R., Dobrev, N., van Emden, T., Hirano, Y., Suma Sreechakram, V.N., Fischer-Burkart, S., Kinugasa, Y., Nevers, A., Rougemaille, M., Sinning, I., Fischer, T., Hiraoka, Y., Braun, S., 2022. The inner nuclear membrane protein Lem2 coordinates RNA degradation at the nuclear periphery. Nat. Struct. Mol. Biol. 29, 910–921. 10.1038/s41594-022-00831-6

McCarthy, R.L., Kaeding, K.E., Keller, S.H., Zhong, Y., Xu, L., Hsieh, A., Hou, Y., Donahue, G., Becker, J.S., Alberto, O., Lim, B., Zaret, K.S., 2021. Diverse heterochromatin-associated proteins repress distinct classes of genes and repetitive elements. Nat. Cell Biol. 23, 905–914. 10.1038/s41556-021-00725-7

Mentch, S.J., Mehrmohamadi, M., Huang, L., Liu, X., Gupta, D., Mattocks, D., Gómez Padilla, P., Ables, G., Bamman, M.M., Thalacker-Mercer, A.E., Nichenametla, S.N., Locasale, J.W., 2015. Histone Methylation Dynamics and Gene Regulation Occur through the Sensing of One-Carbon Metabolism. Cell Metab. 22, 861–873. 10.1016/j.cmet.2015.08.024

Mersman, D.P., Du, H.-N., Fingerman, I.M., South, P.F., Briggs, S.D., 2012. Charge-based interaction conserved within histone H3 lysine 4 (H3K4) methyltransferase complexes is needed for protein stability, histone methylation, and gene expression. J. Biol. Chem. 287, 2652–2665. 10.1074/jbc.M111.280867

Mimura, S., Komata, M., Kishi, T., Shirahige, K., Kamura, T., 2009. SCFDia2 regulates DNA replication forks during S-phase in budding yeast. EMBO J. 28, 3693–3705. 10.1038/em-boj.2009.320

Motamedi, M.R., Hong, E.-J.E., Li, X., Gerber, S., Denison, C., Gygi, S., Moazed, D., 2008. HP1 proteins form distinct complexes and mediate heterochromatic gene silencing by nonoverlapping mechanisms. Mol. Cell 32, 778–790. 10.1016/j.mol-cel.2008.10.026

Motamedi, M.R., Verdel, A., Colmenares, S.U., Gerber, S.A., Gygi, S.P., Moazed, D., 2004. Two RNAi complexes, RITS and RDRC, physically interact and localize to noncoding centromeric RNAs. Cell 119, 789–802. 10.1016/j.cell.2004.11.034

Murphy, P.J., Berger, F., 2023. The chromatin source-sink hypothesis: a shared mode of chromatin-mediated regulations. Dev. Camb. Engl. 150, dev201989. 10.1242/dev.201989

Nakayama, J., Rice, J.C., Strahl, B.D., Allis, C.D., Grewal, S.I., 2001. Role of histone H3 lysine 9 methylation in epigenetic control of heterochromatin assembly. Science 292, 110–113. 10.1126/science.1060118

Nathanailidou, P., Dhakshnamoorthy, J., Xiao, H., Zofall, M., Holla, S., O’Neill, M., Andresson, T., Wheeler, D., Grewal, S.I.S., 2024. Specialized replication of heterochromatin domains ensures self-templated chromatin assembly and epigenetic inheritance. Proc. Natl. Acad. Sci. 121, e2315596121. 10.1073/pnas.2315596121

Nimmo, E.R., Pidoux, A.L., Perry, P.E., Allshire, R.C., 1998. Defective meiosis in telomere-silencing mutants of Schizosaccharomyces pombe. Nature 392, 825–828. 10.1038/33941

Noma, K., Sugiyama, T., Cam, H., Verdel, A., Zofall, M., Jia, S., Moazed, D., Grewal, S.I.S., 2004. RITS acts in cis to promote RNA interference–mediated transcriptional and post-transcriptional silencing. Nat. Genet. 36, 1174–1180. 10.1038/ng1452

Ostermann, K., Lorentz, A., Schmidt, H., 1993. The fission yeast rad22 gene, having a function in mating-type switching and repair of DNA damages, encodes a protein homolog to Rad52 of Saccharomyces cerevisiae. Nucleic Acids Res. 21, 5940–5944. 10.1093/nar/21.25.5940

Oya, E., Nakagawa, R., Yoshimura, Y., Tanaka, M., Nishibuchi, G., Machida, S., Shirai, A., Ekwall, K., Kurumizaka, H., Tagami, H., Nakayama, J., 2019. H3K14 ubiquitylation promotes H3K9 methylation for heterochromatin assembly. EMBO Rep. 20, e48111. 10.15252/embr.201948111

Qu, Q., Takahashi, Y.-H., Yang, Y., Hu, H., Zhang, Y., Brunzelle, J.S., Couture, J.-F., Shilatifard, A., Skiniotis, G., 2018. Structure and Conformational Dynamics of a COMPASS Histone H3K4 Methyl-transferase Complex. Cell 174, 1117–1126.e12. 10.1016/j.cell.2018.07.020

Ragunathan, K., Jih, G., Moazed, D., 2015. Epigenetics. Epigenetic inheritance uncoupled from sequence-specific recruitment. Science 348, 1258699. 10.1126/science.1258699

Reyes-Turcu, F.E., Zhang, K., Zofall, M., Chen, E., Grewal, S.I.S., 2011. Defects in RNA quality control factors reveal RNAi-independent nucleation of heterochromatin. Nat. Struct. Mol. Biol. 18, 1132–1138. 10.1038/nsmb.2122

Roseaulin, L.C., Noguchi, C., Martinez, E., Ziegler, M.A., Toda, T., Noguchi, E., 2013. Coordinated Degradation of Replisome Components Ensures Genome Stability upon Replication Stress in the Absence of the Replication Fork Protection Complex. PLoS Genet. 9, e1003213. 10.1371/journal.pgen.1003213

Rougemaille, M., Braun, S., Coyle, S., Dumesic, P.A., Garcia, J.F., Isaac, R.S., Libri, D., Narlikar, G.J., Madhani, H.D., 2012. Ers1 links HP1 to RNAi. Proc. Natl. Acad. Sci. 109, 11258–11263. 10.1073/pnas.1204947109

Sadeghi, L., Prasad, P., Ekwall, K., Cohen, A., Svensson, J.P., 2015. The Paf1 complex factors Leo1 and Paf1 promote local histone turnover to modulate chromatin states in fission yeast. EMBO Rep. 16, 1673–1687. 10.15252/embr.201541214

Serefidou, M., Venkatasubramani, A.V., Imhof, A., 2019. The Impact of One Carbon Metabolism on Histone Methylation. Front. Genet. 10.

Shimada, A., Dohke, K., Sadaie, M., Shinmyozu, K., Nakayama, J.-I., Urano, T., Murakami, Y., 2009. Phosphorylation of Swi6/HP1 regulates transcriptional gene silencing at heterochromatin. Genes Dev. 23, 18–23. 10.1101/gad.1708009

Shipkovenska, G., Durango, A., Kalocsay, M., Gygi, S.P., Moazed, D., 2020. A conserved RNA degradation complex required for spreading and epigenetic inheritance of heterochromatin. eLife 9, e54341. 10.7554/eLife.54341

Stehling, O., Vashisht, A.A., Mascarenhas, J., Jonsson, Z.O., Sharma, T., Netz, D.J.A., Pierik, A.J., Wohlschlegel, J.A., Lill, R., 2012. MMS19 assembles iron-sulfur proteins required for DNA metabolism and genomic integrity. Science 337, 195–199. 10.1126/science.1219723

Stirpe, A., Guidotti, N., Northall, S.J., Kilic, S., Hainard, A., Vadas, O., Fierz, B., Schalch, T., 2021. SUV39 SET domains mediate cross-talk of heterochromatic histone marks. eLife 10, e62682. 10.7554/eLife.62682

Strachan, J., Leidecker, O., Spanos, C., Le Coz, C., Chapman, E., Arsenijevic, A., Zhang, H., Zhao, N., Spoel, S.H., Bayne, E.H., 2023. SUMOylation regulates Lem2 function in centromere clustering and silencing. J. Cell Sci. 136, jcs260868. 10.1242/jcs.260868

Sugiyama, T., Cam, H., Verdel, A., Moazed, D., Grewal, S.I.S., 2005. RNA-dependent RNA polymerase is an essential component of a self-enforcing loop coupling heterochromatin assembly to siRNA production. Proc. Natl. Acad. Sci. 102, 152–157. 10.1073/pnas.0407641102

Sugiyama, T., Cam, H.P., Sugiyama, R., Noma, K., Zofall, M., Kobayashi, R., Grewal, S.I.S., 2007. SHREC, an effector complex for heterochromatic transcriptional silencing. Cell 128, 491–504. 10.1016/j.cell.2006.12.035

Szklarczyk, D., Kirsch, R., Koutrouli, M., Nastou, K., Mehryary, F., Hachilif, R., Gable, A.L., Fang, T., Doncheva, N.T., Pyysalo, S., Bork, P., Jensen, L.J., von Mering, C., 2023. The STRING database in 2023: protein-protein association networks and functional enrichment analyses for any sequenced genome of interest. Nucleic Acids Res. 51, D638–D646. 10.1093/nar/gkac1000

Tadeo, X., Wang, J., Kallgren, S.P., Liu, J., Reddy, B.D., Qiao, F., Jia, S., 2013. Elimination of shelterin components bypasses RNAi for pericentric heterochromatin assembly. Genes Dev. 27, 2489–2499. 10.1101/gad.226118.113

Taglini, F., Chapman, E., van Nues, R., Theron, E., Bayne, E.H., 2020. Mkt1 is required for RNAi-mediated silencing and establishment of heterochromatin in fission yeast. Nucleic Acids Res. 48, 1239–1253. 10.1093/nar/gkz1157

Takayama, N., Nishimura, S., Nakamura, S., Shimizu, T., Ohnishi, R., Endo, H., Yamaguchi, T., Otsu, M., Nishimura, K., Nakanishi, M., Sawaguchi, A., Nagai, R., Takahashi, K., Yamanaka, S., Nakauchi, H., Eto, K., 2010. Transient activation of c-MYC expression is critical for efficient platelet generation from human induced pluripotent stem cells. J. Exp. Med. 207, 2817–2830. 10.1084/jem.20100844

Taneja, N., Zofall, M., Balachandran, V., Thillainadesan, G., Sugiyama, T., Wheeler, D., Zhou, M., Grewal, S.I.S., 2017. SNF2 Family Protein Fft3 Suppresses Nucleosome Turnover to Promote Epigenetic Inheritance and Proper Replication. Mol. Cell 66, 50–62.e6. 10.1016/j.molcel.2017.02.006

Tange, Y., Chikashige, Y., Takahata, S., Kawakami, K., Higashi, M., Mori, C., Kojidani, T., Hirano, Y., Asakawa, H., Murakami, Y., Haraguchi, T., Hiraoka, Y., 2016. Inner nuclear membrane protein Lem2 augments heterochromatin formation in response to nutritional conditions. Genes Cells Devoted Mol. Cell. Mech. 21, 812– 832. 10.1111/gtc.12385

Tashiro, S., Asano, T., Kanoh, J., Ishikawa, F., 2013. Transcription-induced chromatin association of RNA surveillance factors mediates facultative heterochromatin formation in fission yeast. Genes Cells Devoted Mol. Cell. Mech. 18, 327–339. 10.1111/gtc.12038

Tashiro, S., Handa, T., Matsuda, A., Ban, T., Takigawa, T., Miyasato, K., Ishii, K., Kugou, K., Ohta, K., Hiraoka, Y., Masukata, H., Kanoh, J., 2016. Shugoshin forms a specialized chromatin domain at subtelomeres that regulates transcription and replication timing. Nat. Commun. 7, 10393. 10.1038/ncomms10393

Thakran, P., Pandit, P.A., Datta, S., Kolathur, K.K., Pleiss, J.A., Mishra, S.K., 2018. Sde2 is an intron-specific pre-mRNA splicing regulator activated by ubiquitin-like processing. EMBO J. 37, 89– 101. 10.15252/embj.201796751

Thon, G., Hansen, K.R., Altes, S.P., Sidhu, D., Singh, G., Verhein-Hansen, J., Bonaduce, M.J., Klar, A.J.S., 2005. The Clr7 and Clr8 Directionality Factors and the Pcu4 Cullin Mediate Heterochromatin Formation in the Fission Yeast Schizosaccharomyces pombe. Genetics 171, 1583–1595. 10.1534/genetics.105.048298

Torres-Garcia, S., Di Pompeo, L., Eivers, L., Gaborieau, B., White, S.A., Pidoux, A.L., Kanigowska, P., Yaseen, I., Cai, Y., Allshire, R.C., 2020. SpEDIT: A fast and efficient CRISPR/Cas9 method for fission yeast. Wellcome Open Res. 5, 274. 10.12688/wellcomeopenres.16405.1

Towbin, B.D., González-Aguilera, C., Sack, R., Gaidatzis, D., Kalck, V., Meister, P., Askjaer, P., Gasser, S.M., 2012. Step-wise methylation of histone H3K9 positions heterochromatin at the nuclear periphery. Cell 150, 934–947. 10.1016/j.cell.2012.06.051

Verdel, A., Jia, S., Gerber, S., Sugiyama, T., Gygi, S., Grewal, S.I.S., Moazed, D., 2004. RNAi-mediated targeting of heterochromatin by the RITS complex. Science 303, 672–676. 10.1126/science.1093686

Verrier, L., Taglini, F., Barrales, R.R., Webb, S., Urano, T., Braun, S., Bayne, E.H., 2015. Global regulation of heterochromatin spreading by Leo1. Open Biol. 5, 150045. 10.1098/rsob.150045

Wang, H., Dienemann, C., Stützer, A., Urlaub, H., Cheung, A.C.M., Cramer, P., 2020. Structure of the transcription coactivator SAGA. Nature 577, 717–720. 10.1038/s41586-020-1933-5

Wang, J., Lawry, S.T., Cohen, A.L., Jia, S., 2014a. Chromosome boundary elements and regulation of heterochromatin spreading. Cell. Mol. Life Sci. 71, 4841–4852. 10.1007/s00018-014-1725-x

Wang, J., Tadeo, X., Hou, H., Andrews, S., Moresco, J.J., Yates, J.R., Nagy, P.L., Jia, S., 2014b. Tls1 regulates splicing of shelterin components to control telomeric heterochromatin assembly and telomere length. Nucleic Acids Res. 42, 11419–11432. 10.1093/nar/gku842

Wang, Y., Kallgren, S.P., Reddy, B.D., Kuntz, K., López-Maury, L., Thompson, J., Watt, S., Ma, C., Hou, H., Shi, Y., Yates, J.R., Bähler, J., O’Connell, M.J., Jia, S., 2012. Histone H3 lysine 14 acetylation is required for activation of a DNA damage checkpoint in fission yeast. J. Biol. Chem. 287, 4386–4393. 10.1074/jbc.M111.329417

Yu, R., Wang, X., Moazed, D., 2018. Epigenetic inheritance mediated by coupling of RNAi and histone H3K9 methylation. Nature 558, 615–619. 10.1038/s41586-018-0239-3

Zhang, K., Mosch, K., Fischle, W., Grewal, S.I.S., 2008. Roles of the Clr4 methyltransferase complex in nucleation, spreading and maintenance of heterochromatin. Nat. Struct. Mol. Biol. 15, 381– 388. 10.1038/nsmb.1406

Zofall, M., Grewal, S.I.S., 2007. HULC, a histone H2B ubiquitinating complex, modulates heterochromatin independent of histone methylation in fission yeast. J. Biol. Chem. 282, 14065–14072. 10.1074/jbc.M700292200

Zofall, M., Grewal, S.I.S., 2006. Swi6/HP1 recruits a JmjC domain protein to facilitate transcription of heterochromatic repeats. Mol. Cell 22, 681–692. 10.1016/j.molcel.2006.05.010

Zofall, M., Smith, D.R., Mizuguchi, T., Dhakshnamoorthy, J., Grewal, S.I.S., 2016. Taz1-Shelterin Promotes Facultative Heterochromatin Assembly at Chromosome-Internal Sites Containing Late Replication Origins. Mol. Cell 62, 862–874. 10.1016/j.mol-cel.2016.04.034

Zofall, M., Yamanaka, S., Reyes-Turcu, F.E., Zhang, K., Rubin, C., Grewal, S.I.S., 2012. RNA elimination machinery targeting meiotic mRNAs promotes facultative heterochromatin formation. Science 335, 96–100. 10.1126/science.1211651

